# Lineage-specific regulation and RNA-associated functions of SATB1 in mature B-cells

**DOI:** 10.64898/2026.07.22.739998

**Authors:** Jean-Yves Alejandro Frayssinhes, Lya Meriot, Tiffany Marchiol, Emilie Pinault, Thierry Forné, Charalampos Spilianakis, Eric Pinaud, Sandrine Le Noir

## Abstract

Special AT-rich binding protein 1 (SATB1) is a nuclear matrix-associated transcription factor that orchestrates higher-order chromatin architecture and gene expression in T and B lymphocytes. While SATB1’s role in T cells is well-established, its regulation and function in B cells remain underexplored. Here, we showed that *Satb1* expression in mature B cells is controlled by alternative promoter usage and splicing, with promoters P1 and P3 dynamically switching upon LPS-induced activation, while P2 remains transcriptionally silent. We identified four *Satb1* mRNA isoforms, including two novel transcripts (*Δ9* and *Δ9Δ11*), all of which maintain open reading frames. *Satb1* transcription decreases upon activation while SATB1 protein content evolves differently, suggesting post-transcriptional regulation. SATB1 forms a high-molecular-weight complex in B cells, and co-immunoprecipitation mass spectrometry reveals RNA-binding proteins as its predominant interactors. These findings specify SATB1 expression kinetics during B-cell activation and reveal its role as a regulator of RNA splicing in B cells.

## Introduction

The SATB1 transcription factor was initially discovered through its ability to bind matrix attachment regions (MARs) within the immunoglobulin heavy chain (*IgH*) locus (Dickinson et al., 1992), and was further characterized as a MAR-binding protein (MBP) for its tight association with the nuclear matrix (Alvarez et al., 2000; de Belle et al., 1998; Wang et al., 2010). SATB1 has since emerged as a critical genome organizer that integrates, chromatin looping, transcriptional regulation, and nuclear architecture. SATB1 has been most extensively characterized in T lymphocytes, where it is indispensable for proper thymocyte development (Alvarez et al., 2000; Kakugawa et al., 2017; Kitagawa et al., 2017). SATB1 regulates multiple checkpoints throughout T cell differentiation, from early progenitors to regulatory subsets, by controlling the expression of stage-specific genes including *Rag1* and *Rag2* to promote V(D)J recombination (Hao et al., 2015; Zelenka et al., 2022). SATB1 positively regulates expression of the master regulator *Bcl6*, a critical step for T cell lineage specification, by mediating super-enhancers regions topology and thereby enabling high expression of cell identity genes (Feng et al., 2022; Zelenka et al., 2022). This genome-organizing function is particularly evident in T lymphocytes, where SATB1 orchestrates the three-dimensional configuration of cytokine gene loci and other immune-related genes (Kohwi-Shigematsu et al., 2013; Zelenka and Spilianakis, 2020). Surprisingly, SATB1 also exhibits functional ambivalence, acting as either a transcriptional activator or repressor depending on cellular context. For instance, SATB1 represses *c-myc* in resting T cells but stimulates its expression upon T cell activation (Cai et al., 2003). The recent identification of two SATB1 isoforms, full-length and short (Δ11), displaying different biophysical properties may explain this ambivalence (Zelenka et al., 2023). Despite extensive knowledge of SATB1 in T cell biology, significant gaps remain in understanding its roles in other lineages, particularly in B cells. Initial studies reported modest reductions in B cell numbers in *Satb1* knockout mice (Alvarez et al., 2000). More recent work using fluorescent reporter models revealed dynamic *Satb1* expression throughout B cell differentiation, with elevated expression in naive B cell subsets, and proposed a role for SATB1 in BCR-mediated B cell survival (Ozawa et al., 2022). In contrast to the given SATB1’s established function in regulating *Rag* gene expression in T cells (Hao et al., 2015), our group observed no defect in *Rag* gene expression in early developing B-lineage cells (Thomas et al., 2023). However, the fact that both deletion of MARsEµ and SATB1 impact similarly Igµ and BCR expression in mature B cells strongly support the idea that SATB1 directly regulates immunoglobulin heavy chain transcription (Martin et al., 2023). The functional ambivalence of SATB1 is also observed in mature B cells, in which it represses *IgH* at the resting stage but stimulates its expression upon B cell activation (Thomas et al., 2023).

SATB1 functions as a central chromatin organizer by tethering DNA AT-rich sequences (MARs) from specific genomic loci to the nuclear matrix, thereby establishing tissue-specific chromatin loops interactions (Cai et al., 2006; de Belle et al., 1998; Wang et al., 2023). Several studies support that this anchoring mechanism occurs at specialized subnuclear compartments, such as promyelocytic nuclear bodies (PML-NBs), which belong to the nuclear matrix and organize the major histocompatibility complex (MHC) class 1 locus into distinct higher-order chromatin loop structures (Kumar et al., 2007; Lallemand-Breitenbach and de Thé, 2010; Tan et al., 2008). Through its ability to multimerize (Wang et al., 2014, 2012) and bind multiple DNA elements simultaneously, SATB1 facilitates long-range enhancer-promoter communication and the formation of specialized chromatin compartments (Zelenka et al., 2022). Emerging evidence suggests that SATB1 may also participate in liquid-liquid phase separation, a biophysical mechanism that compartmentalizes nuclear functions and regulates transcription (Feng et al., 2022; Zelenka et al., 2023). SATB1 exhibits remarkable structural and functional conservation between humans and mice, sharing over 98% amino acid identity (Alvarez et al., 2000). The protein contains two CUT domains for DNA binding activity (Nakagomi et al., 1994) and at least two predicted prion-like domain (PrLD) that contribute to RNA binding protein (RBP) function (Seo et al., 2005; Zelenka et al., 2023), in addition to several distinct functional domains that collectively enable its pleiotropic activities. The N-terminal ubiquitin-like domain (ULD) facilitates SATB1 multimerization, enabling homotetramer formation essential for establishing higher-order chromatin structures (Wang et al., 2014, 2012). The CUT-like domains and a homeodomain confer DNA-binding affinity and sequence specificity, respectively (Purbey et al., 2008; Yamasaki et al., 2007). This flexibility is achieved through context-dependent chromatin binding to recognize AT-rich sequences, either associating with euchromatic regions (Cai et al., 2006) or within nucleosome-dense regions (Ghosh et al., 2019; Kohwi et al., 2025), supporting its candidacy as a pioneer transcription factor capable of initiating cell reprogramming (Soufi et al., 2015). The transcriptional activity of SATB1 is dynamically controlled through multiple post-translational modifications, including phosphorylation (Galande et al., 2007; Pavan Kumar et al., 2006), acetylation (Purbey et al., 2009), and SUMO conjugation, which further regulates SATB1 subnuclear localization and stability (Tan et al., 2010, 2008). Furthermore, the *Satb1* gene itself is tightly regulated through alternative promoter usage, a mechanism that fine-tunes mRNA expression, splicing and potentially protein levels during T cell development and differentiation (Khare et al., 2019; Patta et al., 2020). However, SATB1 expression dynamics in B cell biology remain comparatively underexplored.

In this context, we used a conditional *Satb1* knockout mouse model to explore *Satb1* expression dynamics in primary B cells. We found that *Satb1* gene expression is controlled through both alternative promoters switching and alternative splicing of exons 9 and 11. Although *Satb1* transcripts were rapidly downregulated upon LPS stimulation *in vitro*, SATB1 protein levels relatively stable up to 4 days post activation, suggesting that the protein remains functional. Indeed, we readily detected SATB1 associated with RNA-binding proteins by co-IP/MS in both naive and LPS-activated B cells. Together, these multilayered regulatory mechanisms ensure precise control of SATB1 expression and underpin its versatility—from chromatin organization to RNA-associated functions—in normal B-cell physiology.

## Materials and methods

### Mice

To generate a mouse model in which SATB1 is concomitantly suppressed upon BCR activation, we employed the breeding strategy previously described by (Thomas et al., 2023). Briefly, eight-to 10-week-old WT (*Satb1*^flx/+^, *Cd79a*^+/+^) and cHET (*Satb1*^flx/+^, *Cd79a*^cre/+^) mice were bred to generate cKO^SATB1^ (*Satb1*^flx/flx^, *Cd79a*^cre/+^) mice. Mice were maintained in SOPF (Specific and opportunistic pathogen free) conditions at 21-23°C with 12 h light/dark cycle. Procedures were reviewed and approved by the Ministère de l’Enseignement Supérieur, de la Recherche et de l’Innovation, referring to APAFIS #51397-2024101117134127 v3.

### Cell sorting

To sort both resting (B220^+^/GL7^−^) and activated (B220^+^/GL7^+^) Peyer’s patch (PP) B-cells, cells were labelled with anti-B220-APC (RA3-6B2 clone, BD 103212) and GL7-FITC (GL-7 clone, BD 553666) conjugated antibodies. Peyer’s patch B-cells were sorted on ARIA III apparatus (BD Biosciences). To sort splenic resting B-cells, cells were negatively and magnetically sorted using the EasySep™ Mouse B Cell Isolation Kit (STEMCELL, #19854). B-cells population (B220^+^) purity was assessed in flow cytometry using B220 (anti-B220/CD45R-BV711, #103255) and CD43 (anti-CD43-FITC, #553270) cell surface markers and analyzed on CytoFlex apparatus (Beckman Coulter).

### Cell culture and *in vitro* B-cell activation

Splenic B-cells were cultured at 1 × 10^6^ cells/mL in DMEM (1X) – Glutamax medium (Gibco, #32430-027) supplemented with 10% fetal bovine serum (Dutscher, #S1810-500), 2 mM L-glutamine (Eurobio, #CSTGLU00-0U), 1X MEM non-essential amino acids solution (Eurobio, #11140-035), 50 U/mL penicillin‒streptomycin (Gibco, #15070-063), 1 mM sodium pyruvate (Eurobio, #11360-039), and 0,001% 2-β-mercaptoethanol (Sigma‒Aldrich, #60-24-2) in the presence of 1 µg/mL lipopolysaccharide (LPS-Invivogen, #tlrl-3pelps). Primary B-cells were cultured at 37°C and 5% CO_2_. Cells were harvested at indicated time points (resting, and from day 2 to day 5) and processed for further sampling.

### ATAC-sequencing analysis

To analyze chromatin accessibility within *Satb1* promoter regions in splenic (CD4^+^ and CD8^+^ cells) and thymic (CD4 and CD8 single and double positive populations) T-cells, we used ImmGen Datasets (GSE100738). Chromatin accessibility in B cell subsets was evaluated from published datasets (PRJEB59246) (Bruzeau et al., 2024) and in additional samples (this study). In brief, 10 000 resting or 3-day LPS-stimulated B cells were lysed in buffer containing 0.1% Tween-20, 0.1% Nonidet P-40, 0.01% digitonin, 10 mM Tris–HCl pH 7.4, 10 mM NaCl, and 3 mM MgCl₂ for 3 min on ice. Tagmentation was carried out with Illumina transposase in 1× TD buffer supplemented with 0.01% digitonin and 0.1% Tween-20 at 37 °C for 30 min with agitation. Transposed DNA was purified (MinElute kit, Qiagen) and amplified by PCR using the Ad1 and barcoded Ad2.X primers. Libraries were sequenced as 2×100 bp paired-end reads on a NovaSeq 6000 instrument. Raw sequencing reads were trimmed with TrimGalore (retaining only pairs ≥ 55 bp) and processed with the nf-core/atacseq pipeline against the mm10 reference genome. All bioinformatic steps followed the procedures detailed in (Bruzeau et al., 2024). *Satb1* promoter regions accessibility was examined across all experimental conditions with IGV software (v2.19.7).

### RNA-sequencing analysis and Sashimi plots

To analyze SATB1 transcription, we re-analyzed the RNA-sequencing datasets from previous studies in (Thomas et al., 2023) (PRJEB52320) and (Zelenka et al., 2023) (GSE173470). *Satb1* promoter read counts were examined across all experimental conditions with IGV software (v2.19.7). Sashimi plots were generated from BAM files of resting B-cells, LPS-activated B-cells at day2 and in sorted CD138 positives plasmablast cells after 4 days and whole thymus using IGV (v2.19.7). Data are displayed as read per millions (RPM).

### RNA extraction, RT-PCR and quantitative real-time PCR (RT-qPCR)

Total RNA was extracted from splenic resting B cells, LPS-stimulated B cells (from 2 and 5 days of culture), and Peyer’s patch B-cells using TRIzol Reagent (Ambion Life Technologies). RNA purification was performed according to the manufacturer’s instructions. For RT-qPCR analysis, 1 µg of total RNA was treated with DNase I (Invitrogen, #18068-015) to remove residual genomic DNA, and reverse transcription was carried out using the High-Capacity cDNA Reverse Transcription Kit (Applied Biosystems, #4368814) following the recommended protocol. RT-PCR of *Satb1* promoter to exon 4 was performed with the designed primers (P1F, P2F, P3F, P4F and R4), represented **in Figure S1.A**, using 1 ng/µL equivalent reverse transcribed RNA template (cDNA) from three splenic resting B cells independant cDNA template. Each PCR reaction contained 10 μL of Taq’Ozyme OneMix w/Dye (2X) (OZYME, #OZYA011-1000), 10 μM of each primer (forward and reverse) completed with H_2_O (Aguettant, #69802A) in 20 μL final. The following cycling conditions were used: 98°C for 3 min; 35 cycles (98°C for 30 sec, 62°C for 45 sec, 72°C for 1 min); 72°C for 5 min. Products were resolved on a 2.2% agarose gel **(Figure S2.B)**.

RT-PCR *Satb1* exon 12-7 and *Satb1* exon 12-8 were performed with 1 ng/µL equivalent reverse transcribed RNA template from splenic resting B cells. Each PCR reaction contained 0.8 μL of LongAmp® Taq DNA Polymerase (NEB, #M0323L), 10 μM of each primer (forward and reverse), 0.6 μL of 10 mM dNTPs (Invitrogen, #NTPMX100C), 4 μL of 5 × LongAmp Taq Reaction Buffer (NEB, #B0323S) completed with H_2_O (Aguettant, #69802A) in 20 μL final. For the *Satb1* exon 12-7 RT-PCR, the following cycling conditions were used: 94°C for 30 sec; 35 cycles (94°C for 30 sec, 62.1°C for 15 sec, 65°C for 25 sec); 65°C for 2 min 30 sec. PCR products were resolved in 1.5% agarose gel. For the *Satb1* exon 12-8 RT-PCR, the following cycling conditions were used: 94°C for 30 sec; 35 cycles (94°C for 30 sec, 62.1°C for 15 sec, 65°C for 25 sec); 65°C for 2 min 30 sec. PCR products were resolved in 2% agarose gel. All primer sequences for RT-PCR are listed in Supplementary Table S1.

RT-PCR of *Satb1* spliced isoform was performed with the designed primers (Full, R11-10 and F9-10; Δ11, R12-10 and F9-10; Δ9, R11-10 and F8-10; and Δ9Δ11, R12-10 and F8-10) using 1 ng/µL equivalent reverse transcribed RNA template (cDNA) or genomic DNA (gDNA) from splenic resting B cells as template. Each PCR reaction contained 10 μL of Taq’Ozyme OneMix w/Dye (2X) (OZYME, #OZYA011-1000), 10 μM of each primer (forward and reverse) completed with H_2_O (Aguettant, #69802A) in 20 μL final. The following cycling conditions were used: 98°C for 1 min; 35 cycles (98°C for 30 sec, 62°C for 45 sec, 72°C for 20 sec); 72°C for 5 min. Products were resolved on a 3% agarose gel. Quantitative PCR was performed on 20 ng of equivalent reverse transcribed RNA using the SensiFAST SYBR Hi-ROX kit (BioLine, #BIO-92020) on a QuantStudio 3 system (Applied Biosystems). *Satb1* promoter transcript levels were quantified with promoter-specific primers designed according to (Patta et al., 2020). *Satb1* spliced isoforms transcript levels were quantified with isoform-specific primers designed with Genome Data Viewer (NCBI). The RT-qPCR run was programmed as following: stage 1 (95°C for 2 min), stage 2 (95°C for 10 sec), stage 3 (60°C for 30 sec). Stages 2 and 3 were repeated to 40 cycles. Melting curve analysis was programmed as following: stage 1 (95°C for 15 sec), stage 2 (60°C for 1 min), stage 3 (95°C for 15 sec). Relative quantification (RQ) indicates a gene expression level relative to a control sample. The formula used was RQ = 2^−ΔΔCt^, where ΔCt is the difference between the Ct value of the target gene and the Ct value of the reference gene within the same sample. ΔΔCt is then calculated by subtracting the ΔCt of the control from the ΔCt of the experimental sample. Expression was normalized to either *Gapdh* or *Actb* as endogenous control genes. Fold change was calculated compared to either resting splenic or Peyer’s patch (B220^+^/Gl7^−^) B-cells. Multiple T_m_ peaks during melting curve analysis were assessed to verify nonspecific amplification, specifically for isoform-specific primers in RT-qPCR. All primer sequences for RT-qPCR are listed in Supplementary Table S1.

**Supplementary Table S1:**
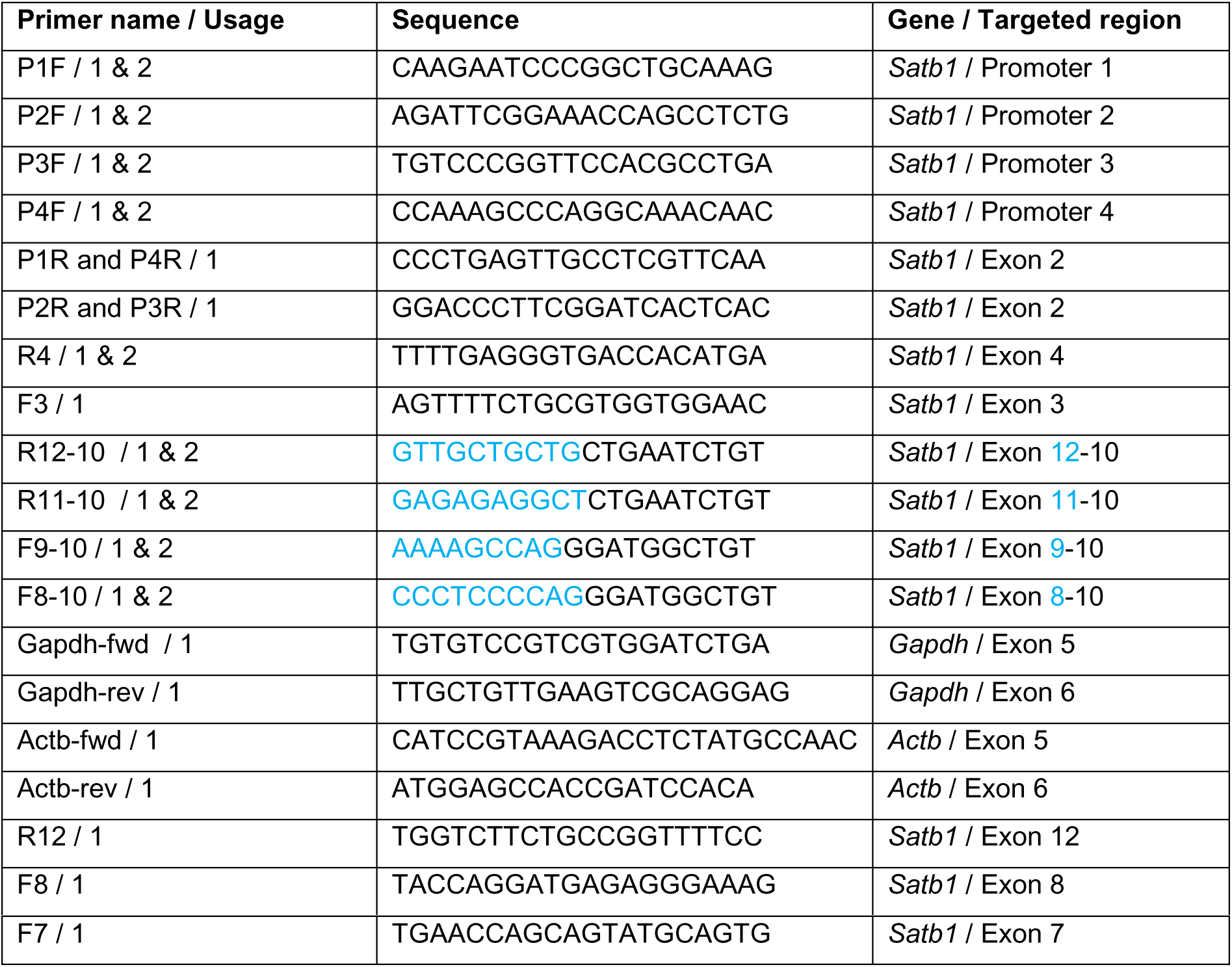
summary table of primers used in this study specifically designed in *mus musculus* for RT-qPCR and RT-PCR experiments. Usage: 1, indicates usage for RT-qPCR; 2, indicates usage for RT-PCR. Sequence highlighted in blue color indicates the overlapped exon for custom-primers.

### gDNA extraction and RT-PCR

Genomic DNA (gDNA) from splenic resting B-cells were prepared as following: freshly sorted-resting B-cells were washed twice in PBS (1×)., and then were incubated in gDNA lysis buffer (NaCl 200 mM, HEPES 10 mM pH 8, EDTA 2 mM, SDS 0.2%, Proteinase K 0.2 µg/µL), overnight at 56°C. Then, gDNA was further precipitated in 500 µL Isopropanol, washed twice in 1 mL ethanol (70 %), and then air dried. DNA pellet was resuspended in H_2_O (Aguettant, #69802A).

### Cellular fractionation and protein extraction

Cells were harvested and washed twice with PBS (1×). For 5 × 10⁵ cells, the cell pellet was resuspended in 100 µL of nuclei extraction buffer containing glycerol (20%), HEPES (20 mM, pH 8.0), KCl (10 mM), Triton X-100 (0.1%), and MnCl₂ (1 mM), supplemented immediately before use with 1× Protease Inhibitor Cocktail (Roche) and 0.5 mM spermidine. Cells were incubated on ice for 10 min. Nuclei were then pelleted by centrifugation at 600 × g for 3 min at 4°C. The supernatant (cytoplasmic fraction) was collected separately, and the pellet represented the isolated nuclear fraction. All steps were performed and adapted from the protocols described in (Lecouvreur et al., 2025; Rebouissou et al., 2022).

To extract proteins from either isolated nuclei or whole cells, a pre-lysis buffer containing HEPES (50 mM, pH 8.0), NaCl (150 mM), Nonidet P-40 (1%), sodium deoxycholate (0.5%), and SDS (0.1%) was prepared and stored at –20°C. Immediately before use, the pre-lysis buffer was supplemented with sodium orthovanadate (1 mM, pH 9.7), PMSF (0.5 mM), and Protease Inhibitor Cocktail (1 mM, Sigma-Aldrich). Nuclei or cells were resuspended in an appropriate volume of lysis buffer on ice (20–80 µL for 1–30 × 10⁶ cells) and immediately centrifuged at 14,000 × g for 10 min at 4°C. The supernatant (nuclear or whole-cell lysate) was collected and stored at –70°C. Protein lysates could be further treated with benzonase to study SATB1 complex in native condition (see below). Protein lysates were quantified using Pierce™ BCA Protein Assay Kits (Thermo Scientific, #23225) according to the manufacturer’s instructions. All reagents are listed in Supplementary Table S2.

For benzonase treatment, nuclear lysates (3 µg/µL) were incubated with benzonase (5 U/µL) in lysis buffer supplemented with 1 mM MgCl₂ for 20 min at 25°C. An aliquot containing 10 µg of benzonase-treated lysate (from two independant WT and cKO^SATB1^ resting B-cells) was then resolved by native-PAGE (gradient 4-10%).

**Supplementary Table S2:** summary table of material for nuclei isolation, cell lysis and protein extraction performed in this study.

| Material | Manufacturer | Reference |
| --- | --- | --- |
| Glycerol | Sigma-Aldrich | G6279-500ML |
| HEPES | Sigma-Aldrich | H3375-100G |
| KCl | Sigma-Aldrich | 1.04936.1000 |
| Triton X-100 | Sigma-Aldrich | T9284 |
| MnCl<sub>2</sub> | Sigma-Aldrich | 85558-1G |
| Protease Inhibitor Cocktail | Roche Diagnostics | 11873580001 |
| Spermidine | Sigma-Aldrich | 85558-1G |
| Sodium deoxycholate | Sigma-Aldrich | 30970-100G |
| Sodium orthovanadate | Sigma-Aldrich | S-6508 |
| NaCl | Sigma-Aldrich | S7653-5KG |
| Nonidet P-40 Substitute | Sigma-Aldrich | 74385-1L |
| Protease Inhibitor Cocktail | Sigma-Aldrich | P8340 |
| PMSF | Sigma-Aldrich | P-7626-25G |
| Benzonase nuclease HC<br>(250U/μl) | Merckmillipore | 71206 |
| SDS | Fishersci | BP166-5 |

### Denaturing SDS-PAGE and Western blot

Whole-cell lysates (10 µg in 20 µL final volume) from resting or LPS-activated B-cells (+1 to +4 days) were prepared in 1× Laemmli buffer and denatured at 95°C for 5 min in a MasterCycler nexus x2. Denatured samples were resolved by SDS-PAGE using 12% TGX Stain-Free FastCast Acrylamide gels (1 mm spacer, 10-well; Bio-Rad, #1610185). Precision Plus Protein Dual Color Standards (Bio-Rad, #161-0374) and Precision Plus Protein All Blue Prestained Protein Standards (Bio-Rad, #161-0373) were used as molecular weight markers (3 µL per well). Electrophoresis was performed in SDS-migration buffer containing Tris-HCl (25 mM), glycine (192 mM), and SDS (0.1%), pH 8.3. The running program was set at 90 V, followed by 120 V for 2 h at room temperature.

Protein loading was assessed using stain-free technology prior to transfer (2 min 30 s activation; 4 s exposure). PVDF Immobilon membranes (Millipore, #ISEQ85R) were activated with ethanol (90%) for 10 min and equilibrated in 1× transfer buffer for at least 15 min at room temperature. Semi-dry transfer was performed using the Trans-Blot Turbo Transfer System (Bio-Rad) at 25 V and 1.3 A for 15 min. After transfer, membranes were blocked in blocking buffer containing Tris (20 mM), glycine (107 mM), pH 7.5, and 5% (w/v) non-fat dry milk (Regilait) for at least 1 h at 4°C with gentle agitation (tube roller).

Membranes were incubated overnight at 4°C with primary antibody (anti-SATB1, 1:5,000, ab109122, Abcam) in blocking buffer. Membranes were then washed three times with PBS-T (PBS containing 0.1% Tween-20; 50 mL each; first two washes for 10 s, last wash for at least 1 h at 4°C). Membranes were incubated with HRP-conjugated secondary antibody (1:5,000, #1706515, Bio-Rad) in blocking buffer for 1 h at 4°C, followed by washing as described above. Immunodetection was performed using either Pierce ECL Western Blotting Substrate (Thermo Fisher, #32106) or high-sensitivity chemiluminescent substrate (Immobilon ECL Ultra, #WBULS0100). All blots were visualized using the Fusion FX6 imaging system (Vilber). Laemmli buffer recipes for SDS-PAGE or native-PAGE were adapted from previous work (Frayssinhes et al., 2021; Jonik-Nowak et al., 2018) and are detailed in **Supplementary Table S3**.

### Native-PAGE

For native condition, whole-cell lysates were resolved using commercial blue native PAGE (BN-PAGE, 4–16% gradient, Invitrogen, #BN2111BX10) according to the manufacturer’s instructions. NativeMark Unstained Protein Standard (Invitrogen, #LC0725) was used as molecular weight marker (3 µL per well). Alternatively, whole-cell lysates or benzonase-treated nuclear lysates were resolved using homemade 4–16% gradient native-PAGE.

Homemade gradient native-PAGE (4–16%) was prepared as follows. Two solutions of acrylamide (4% and 16%) were prepared separately, both without SDS. Immediately before casting, TEMED and ammonium persulfate (APS) were added to both solutions and mixed thoroughly. Gradient casting was performed by aspirating the 4% solution into a 5 mL serological pipette, followed by the 16% solution. Prior to reaching the last 0.5 mL of the 16% solution, an air bubble (approximately 0.5 mL) was intentionally aspirated, followed by the remaining 0.5 mL of the 16% solution. The pipette was gently flicked to allow the air bubble to rise through both solutions, generating the gradient. The mixed solution was then slowly poured into the glass cassette. For casting a gel with a 1 mm spacer, a stoichiometric volume of solution 4% and 16% (3.2 mL : 3.2 mL) was used. After polymerization and sample loading, electrophoresis was performed at 150 V for 1.5 h at room temperature in a migration buffer identical to SDS-migration buffer but without SDS. Homemade gradient 4–10% or 4–16% SDS-PAGE was performed similarly but with SDS included. Detailed compositions are provided in **Supplementary Table S4**.

**Supplementary Table S3:** composition of Laemmli buffer (5×) and native-loading buffer. To prepare 1 mL of 5× loading buffer, add 0.5 mL Glycerol (100%), 125 µL SDS (20%), 125 µL Tris (0.5 M, pH 6.8), 100 µL H_2_O milli-Q (≥ 18 MΩ.cm), 50 µL β-mercaptoethanol (100%). Use the Eppendorf tube graduations to measure 0.5 mL of pure glycerol. Native-loading buffer was prepared identically, except β-mercaptoethanol and SDS were replaced with Milli-Q H_2_O.

| Component | Stock concentration | Final concentration | Reference |
| --- | --- | --- | --- |
| Glycerol | 100% | 50% | G6279-500ML |
| SDS | 20% | 2.5% | BP166-5 |
| Tris-HCl | 0.5 M, pH 6.8 | 62.5 mM | T6066-1KG |
| β-mercaptoethanol | 100% | 0.5% | 60-24-2 |
| Bromophenol | 0.5% | 0.05% | B-8026 |

**Supplementary Table S4:** composition of homemade gradient native or SDS-PAGE gels (4–10% or 4–16%). For a single gradient native gel, replace SDS with Milli-Q H_2_O. Final concentrations are as follows: 0.375 M Tris-HCl (pH 8.8), 0.1% SDS (for denaturing gels), 0.05% APS, 0.1% TEMED, and the desired acrylamide percentage (custom-chosen). Acrylamide/Bis (37.5:1, 30%) was from Euromedex (#EU0088-C).

| Component | Solution (4%) | Solution (10%) | Solution (16%) |
| --- | --- | --- | --- |
| Milli-Q H<sub>2</sub>O | 1.94 mL | 1.30 mL | 0.66 mL |
| Tris-HCl (1.5 M, pH 8.8) | 0.8 mL | 0.8 mL | 0.8 mL |
| SDS (20%) if SDS-PAGE | 16 µL | 16 µL | 16 µL |
| Acrylamide/Bis 37,5:1 (30%) | 0.43 mL | 1.07 mL | 1.71 mL |
| APS (10%) | 16 μL | 16 μL | 16 μL |
| TEMED | 3.2 μL | 3.2 μL | 3.2 μL |

### Two-dimensional gel electrophoresis (native- vs SDS-PAGE)

For two-dimensional electrophoresis, proteins were first separated under native conditions in the first dimension. Benzonase-treated nuclear lysates (10 µg) from WT or cKO^SATB1^ resting B-cells were loaded onto a homemade 4–16% native-PAGE gradient gel, as described above. A lane containing native molecular weight markers was also loaded to verify migration. Following electrophoresis, the entire lane of interest, including the marker lane, was excised vertically from the native gel using a clean scalpel. The excised gel strips were wrapped in cellophane and stored at –20°C until further use. The marker lane was stained with Serva blue staining to control the migration.

For the second dimension (denaturing SDS-PAGE), the stored gel strips were rehydrated in Milli-Q H₂O for 20 min at room temperature with gentle agitation. After removing the water, 250 µL of 2× Laemmli buffer were added to the wrapped strips, which were then incubated for 30 min at 4°C with gentle agitation. The denatured gel strips were then transferred horizontally onto the stacking gel of a second-dimension SDS-PAGE gel (4% stacking, 12% running; either homemade casted or using 12% TGX Stain-Free FastCast Acrylamide gels). SDS-PAGE and subsequent Western blotting were performed as described above. Following transfer to PVDF membrane, the gel was either stained with Serva Blue to confirm complete protein transfer, and the membrane was probed with anti-SATB1 antibody (1:5,000, ab109122, Abcam). Alternatively, stain-free gels were visualized prior transfer to PVDF membrane to control the 1^st^ dimension migration.

### SATB1 co-immunoprecipitation

For SATB1 co-immunoprecipitation (co-IP), nuclear extracts were isolated from resting or LPS-activated splenic B-cells (+2 days) from wild-type (WT) or cKO^SATB1^ mice (aged 2–4 months) and pooled. For each SATB1 pull-down, 9 steps were performed, Step 1: 150 µg of nuclear extract was brought to a volume of 120 µL with buffer A (150 mM NaCl, 50 mM HEPES pH 8.0, 5% glycerol, 0.05% Nonidet P-40, 1 mM PMSF). To pre-clear the nuclear lysate, 20 µL of Dynabeads® protein A (30 mg/mL, Invitrogen, #10001D) were added and incubated overnight at 4°C with gentle agitation (MOCK), in a 50 mL falcon and on a tube roller. Step 2: the supernatant (pre-cleared material) was then transferred to a fresh Eppendorf tube containing 10 µL of Dynabeads® protein A pre-coated with 50 ng of either the monoclonal mouse anti-SATB1 antibody (Santa Cruz, #sc-376096) or the polyclonal rabbit anti-full-length SATB1 antibody (Zelenka et al., 2023). The mixture was incubated overnight at 4°C with gentle agitation. Buffer B was identical to buffer A but without Nonidet P-40. All remaining steps from 3 to 9 were adapted from (Zelenka et al., 2023) and are illustrated in **Figure S6**. Also, all fractions (#) were collected following IP, for SATB1-immunoblotting (IB) to identify in which fraction SATB1 was detected. Input (fraction 1) corresponds to 10 µg of nuclear lysate. Post-IP (fraction 2) is the 150 µL remaining supernatant after SATB1 pull-down. The beads were washed twice with 30 µL of buffer A (fractions 3 and 4), then twice with 30 µL of buffer B (fractions 5 and 6). Bound proteins were eluted twice with 8 M urea (20 µL then 15 µL) for 10 min on ice (fractions 7 and 8). After elution, the beads were resuspended in 20 µL of 1× Laemmli buffer (fraction 9). MOCK beads were processed starting from step 3 in parallel through the same washing and elution steps to generate the corresponding MOCK fractions. All fractions were collected and analyzed by SDS-PAGE and immunoblotting. For Western blot analysis, following fractions were prepared: input (fraction 1, 10 µg), Post-IP (fraction 2, 10 µL), Washing 1 (fraction 3, 10 µL), Washing 2 (fraction 4, 10 µL), Washing 3 (fraction 5, 10 µL), Washing 4 (fraction 6, 10 µL), Elution 1 (fraction 7, 5 µL), Elution 2 (fraction 8, 5 µL), and Beads from IP-SATB1 (fraction 9, 20 µL) or MOCK (fraction 9.M, 20 µL). Collected fractions were resolved by SDS-PAGE on 12% TGX Stain-Free FastCast acrylamide gels (Bio-Rad), as described above. The remaining 15 µL of fractions 7 and 7.M were subjected to mass spectrometry analysis.

### NanoLC-MS/MS analysis

For protein digestion, eluted fractions (20 µL in 8 M urea) were reduced by adding 2 µL of 50 mM dithiothreitol (DTT) and incubated at 56°C for 15 min. Proteins were then alkylated with 2 µL of 100 mM iodoacetamide for 15 min in the dark. Samples were diluted with 4 µL of 1 M ammonium bicarbonate and 124 µL of H₂O to reduce the urea concentration to ≤1 M. CaCl₂ was added to a final concentration of 0.5 mM as a cofactor for thermolysin. Digestion was performed overnight at 37°C after adding 10 µL of thermolysin (0.2 µg/µL). Following digestion, peptides were purified using solid-phase extraction on an Oasis HLB 1 cc (30 mg) cartridge (WAT094225). The cartridge was conditioned sequentially with 1 mL methanol and 1 mL H₂O containing 0.5% formic acid (FA). The digested sample was loaded with 750 µL H₂O/0.5% FA, washed with 1 mL H₂O/0.5% FA, and dried for 5 min. Peptides were eluted with 1 mL methanol and evaporated to dryness under nitrogen flow at 35°C. Dried peptides were reconstituted in 30 µL of mobile phase A (2% acetonitrile [ACN] / 0.1% FA), filtered through a 0.22 µm filter, and analyzed by nanoLC-MS/MS. Peptides were separated using a nanoElute® 2 liquid chromatography system (Bruker) coupled to a timsTOF Pro 2 mass spectrometer (Bruker) operated in DDA-PASEF mode. Samples (5 µL injection) were loaded onto a PepSep Twenty-five column (150 µm × 25 cm, 1.5 µm, 100 Å; Bruker #1893476) maintained at 50°C. The flow rate was set to 400 nL/min using mobile phase A (2% ACN / 0.1% FA) and mobile phase B (100% ACN / 0.1% FA). Peptides were eluted with a gradient from 2% to 25% B over 30 min, with a total run time of 50 min. Mass spectrometry was performed in positive ion mode with a standard 1.1-sec cycle time. TIMS settings were: 0.6 to 1.6 V·s/cm² with a 100-ms accumulation time. Precursor ions with charge ≥2 were selected for fragmentation. MS1 spectra were acquired over m/z 100–1700 with a 100-ms accumulation time. PASEF MS/MS consisted of 10 cycles of 25 MS/MS events (100-ms accumulation each). Samples were injected in the following order: bead-only controls, then complexes, with knockout samples analyzed before wild-type samples within each group. Raw MS/MS data were processed using Mascot (Matrix Science) against the SwissProt database (Mus musculus, 2024-03). Note that the SATB1 entry in SwissProt (SATB1_MOUSE) corresponds to the Δ11 isoform. Therefore, when SATB1 was identified, an additional search was performed using custom sequences including all SATB1 isoforms of interest.

### SATB1 interactome analysis

Proteins were identified using Mascot (Matrix Science) against the SwissProt database (Mus musculus, 2024-03). Search parameters were as follows: peptide mass tolerance of ±20 ppm, fragment mass tolerance of ±0.05 Da, enzyme specificity set to thermolysin (cleavage before Leu, Ile, Phe, Val, Ala; after Leu, Ile, Phe, Val, Ala), with a maximum of two missed cleavages. Only proteins identified with at least two unique peptides and a Mascot score ≥ 30 were considered for further analysis. Proteins uniquely identified in WT SATB1 IP samples or significantly enriched compared to controls were retained as high-confidence SATB1 interactors. For clustering and network analysis, the filtered list of high-confidence interactors was submitted to the STRING database to retrieve protein-protein interaction data. Network nodes were clustered using k-means clustering with vector representations (Szklarczyk et al., 2023). All analyses were performed with n = 3 biological replicates for WT and *n*= 2 for cKO^SATB1^ from resting and + days 2 LPS-activated B-cells.

### Box-plot representations

Whisker plots showed the lower and upper percentile, representing observations outside the 1–99 percentile range. The diagram also shows the median (line), mean (cross), and outlier observations.

### Statistical analysis

Two-tailed Mann–Whitney U tests were used for statistical analysis using GraphPad Prism software: P > 0.05 (ns), P < 0.05 (*), P < 0.01 (**), P < 0.001 (***) and P < 0.0001 (****).

## Results

### Dynamics of *Satb1* promoter activity in mature B-cells

Two studies characterized at least four distinct promoters within the 5′UTR of the *Satb1* gene and provided evidence for an alternative promoter switch of *Satb1* transcription from naive to activated T lymphocytes (Khare et al., 2019; Patta et al., 2020). However, this dynamic has not been investigated in the context of B-cell activation. To explore *Satb1* promoter usage in wild-type (WT) B cells, we first investigated changes in chromatin accessibility within the promoter region. We used publicly available ATAC-seq data from T-cells (ImmGen Datasets, GSE100738) and compared it to our ATAC-seq data from WT resting and LPS-activated B-cells (+3 days). As expected, we observed that all promoter sites were in an open configuration in T-cells **(Figure 1.A, top panel)**. In contrast, the P2 promoter site remained in a closed conformation in resting and LPS-activated B-cells (+3 days), suggesting that *Satb1* transcription does not occur from this specific site **(Figure 1.A, bottom panel)**. To confirm that the P2 promoter site does not promote *Satb1* transcription, we investigated the *Satb1* transcription from each promoters sites in resting and LPS-activated B-cells (day 2) and in plasmablast CD138+ sorted after 4 days of LPS stimulation from our RNA-seq datasets (Thomas et al., 2023). As expected, we observed no active transcription originating from P2, neither in naive nor LPS-activated B-cells, confirming that the transcriptional inactivity of P2 correlates with its closed chromatin state in both conditions. Instead, we mainly observed transcription that occurs from P1, P3 and P4 promoter sites. In addition, all promoter transcription was downregulated upon LPS-induced B-cell activation (10-fold less read counts in + day 4 LPS-activated B-cells compared to the resting state). Moreover, *Satb1* promoter usage shifted from P3 to P1 in CD138 positive B cells **(Figure 1.B)**. To investigate *Satb1* promoter dynamics in B-cells subsets, we quantified the transcriptional output of each promoter by quantitative and standard RT-PCR using primers from a previous study (Patta et al., 2020), schematically represented in **Figure S1.A**. First, to verify all promoter activity in splenic resting B-cells, we tested by RT-PCR whether *Satb1* transcription could reach *Satb1* exon 4, regardless of the promoter site origin. We successfully amplified forward *Satb1* cDNA from P1, P3, and P4, and we also confirmed the absence of forward *Satb1* transcription from P2 **(Figure S1.B)**. Second, we assessed the dynamics of *Satb1* promoter transcription in splenic resting and LPS-activated B-cells (over 5 days) and in naive (B220^+^/GL7^−^) and activated (B220^+^/GL7^+^) Peyer’s patch (PP) B-cells. Quantification of relative promoter activity revealed a decrease in transcripts originating from all promoter sites upon LPS activation *in vitro* **(Figure 1.C, left)**. A similar decrease was also observed *in vivo* when comparing naive to activated B-cells from Peyer’s patches **(Figure 1.C, right)**. Furthermore, we confirmed the absence of active transcription from P2, in resting and activated B-cells both *in vitro* and *in vivo* **(Figure 1.C)**. We further analyzed the proportions of all promoter usage at each indicated time point (from 0 to 5 days post-LPS stimulation) relative to a total of 100%. In splenic resting and naive Peyer’s patch B-cells, P3 is the predominant promoter among the four. Overall, *Satb1* promoter activity decreased with activation. However, the predominant promoter gradually shifted from P3 to P1 as cells transitioned from a resting to an activated stage *in vitro* **(Figure 1.D)**, in agreement with our RNA-seq data. We then compared resting to LPS-activated B-cells at day 5. The P3 promoter contribution decreased from 54% to 34%, while P1 increased from 41% to 64% **(Figure 1.D, left)**. Similar to *in vitro* activated B-cells, *Satb1* promoter activity drastically decreased in activated Peyer’s patch B-cells. Unlike LPS-activated B-cells, activated Peyer’s patch B-cells did not show such a drastic shift in promoter preference, but P3 remained the principal active promoter (although a minor increase in P1 and P4 was observed) **(Figure 1.D, right)**. Importantly, the *Satb1* downregulation observed by RT-qPCR could be a consequence of fluctuating housekeeping gene expression upon B-cell stimulation. We therefore quantified *Actb* transcript levels in addition to *Gapdh* and found similar *Satb1* promoter expression dynamics **(Figure S1.C and S1.D)**. Our data *in vitro* and *in vivo* indicated that P4 contribution was minor in B-cells, reaching at maximum 15% in activated Peyer’s patch B-cells **(Figure 1.D)**, but remained relatively steady, similar to that observed in T-cells (Patta et al., 2020). We hypothesized that, in *mus musculus*, P4 transcription might occur from the reverse strand, which has been previously described in *Homo sapiens* (*SATB1*-*AS1*, *Satb1* antisense long non-coding RNA) (Wang et al., 2025). To explore this possibility, we analyzed total read counts from our RNA-seq data and compared forward versus reverse strands. While P1, P3 and P4 showed classical bidirectional transcription activity, P2 remained inactive in both directions. Indeed, P4 displayed significant antisense transcription on the reverse strand in our B-cells datasets **(Figure 1.E)**. Strikingly, the P4 promoter region is also the initiation site of reverse transcription in T-cells and displayed, in both lymphocyte lineages, splicing patterns towards an upstream region **(Figure S1.E)**.

**Figure 1.**
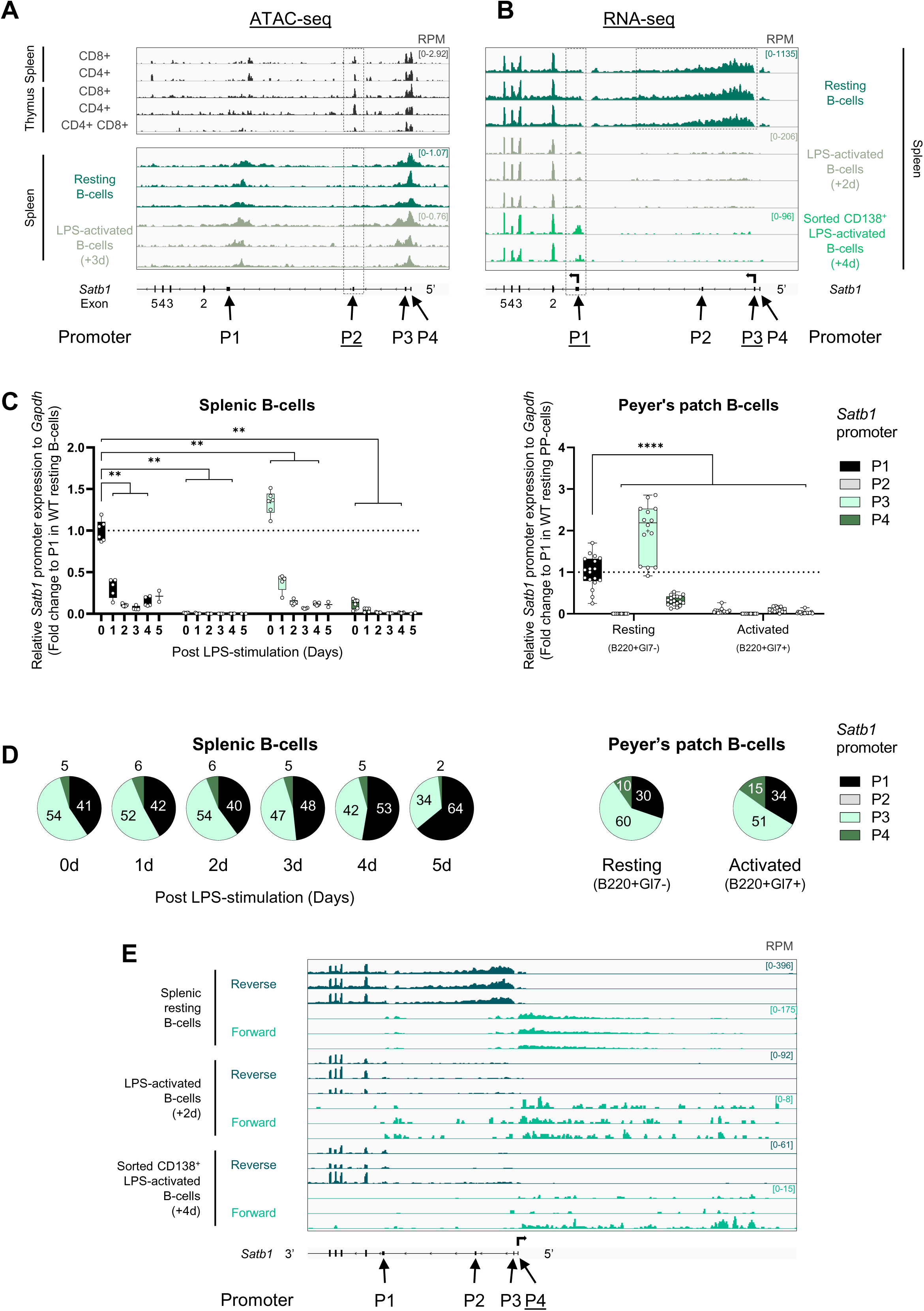
Dynamic of *Satb1* promoter activity in mature B-cells. **(A)** Chromatin accessibility at *Satb1* promoter regions in wild-type (WT) splenic resting and LPS-activated B-cells (+3 days) **(bottom panel)** and in T-cells **(top panel)**, using publicly available ATAC-seq data from T-cells (ImmGen Datasets, GSE100738). **(B)** RNA-seq analysis of *Satb1* promoter usage in WT splenic resting, LPS-activated B-cells (day2) and in plasmablast CD138 positives cells, sorted after 4 days of LPS stimulation. **(C)** Whisker plots showing quantitative RT-PCR analysis of *Satb1* promoters in splenic and Peyer’s patch (PP) B-cells from WT mice. Splenic B-cells **(right panel)** were cultured with LPS for up to 5 days. Expression was normalized to *Gapdh*, and fold change was calculated relative to P1 promoter usage in resting B-cells. *n* = 6 for resting and LPS-activated days 1–4; *n* = 2 for LPS-activated day 5. Peyer’s patch B-cells **(left panel)** were sorted into two populations: resting (B220⁺/GL7⁻) and activated (B220⁺/GL7⁺). Normalization and fold change were performed as in the right panel, with resting Peyer’s patch B-cells as the reference, *N* = 18. **(D)** Pie charts representing the quantitative RT-PCR data from **(C)** for each *Satb1* alternative promoter in splenic B-cells **(left panel)** and Peyer’s patch B-cells **(right panel)** relative to a total of 100%. **(E)** Analysis of three RNA-seq datasets (forward and reverse strands) from WT splenic resting, LPS-activated B-cells (day2) and in plasmablast CD138 positives cells, sorted after 4 days of LPS stimulation. P-value was determined using a two-tailed Mann–Whitney U test; only significant differences are indicated: P < 0.01 (**); P < 0.0001 (****).

### Divergent kinetics of *Satb1* transcript and SATB1 protein levels in B-cells

To quantify total *Satb1* mRNA transcripts, we used primers that specifically amplify *Satb1* exons 3 to 4, a region conserved across all known isoforms (**Figure S2A**) (Zelenka et al., 2023). *Satb1* expression was downregulated both *in vitro* upon LPS-stimulation and *in vivo* in activated Peyer’s patch B-cells, compared to resting B-cells (normalized to *Gapdh)*, and was absent in the *Satb1* conditional knockout model (cKO^SATB1^), which specifically lacks exon 4 in developing B-cells **(Figures 2.A and B and S2A)** (Thomas et al., 2023). Total *Satb1* transcript levels dropped rapidly, on average 10-fold below basal expression after 2 days and up to 5 days of LPS stimulation, or in activated Peyer’s patch B-cells, in agreement with our RNA-seq data **(Figure S2.B and C)**. After just 1 day of LPS activation, 69% of *Satb1* transcripts were downregulated relative to the resting state when normalized to *Gapdh* **(Figure 2.A and 2.B)**. Similar kinetic was observed when normalizing to *Actb*, showing an 80% decrease at day 1 **(Figures S2.B and S2.C)**. Such a drastic drop in *Satb1* transcription suggests tight control of its mRNA levels. To quantify SATB1 protein levels in B-cells, we used a commercial antibody specifically targeting the C-terminal region of SATB1 for immunoblotting **(Figure S2.D)**. Since SATB1 exerts nuclear functions in T-cells, we assumed SATB1 is a nuclear protein in primary B-cells. To verify its subcellular localization, nuclear and cytosolic extracts were prepared from WT and cKO^SATB1^ resting and LPS-activated B-cells following a cell fractionation protocol (Lecouvreur et al., 2025; Rebouissou et al., 2022). As expected, SATB1 was exclusively detected in the nuclear fraction and migrated as a single band on a homemade 4–10% gradient SDS-PAGE **(Figure 2.C)**. To investigate SATB1 protein dynamics in B-cells, cells were collected up to 4 days after LPS stimulation and lysed. Whole-cell lysates (WCL) from WT and cKO^SATB1^ B-cells were used for Western blotting. Total SATB1 protein levels were normalized to GAPDH and fold change was calculated to WT resting B-cells. As expected in our negative control, SATB1 protein was undetectable in cKO^SATB1^ B-cells **(Figures 2.D** and **S2.E)**. In the WT context, SATB1 protein levels showed a transient increase at day 1, followed by a rapid decrease during *in vitro* B-cell activation **(Figure 2.E)**. This contrasts sharply with our total *Satb1* transcript quantification. Indeed, we found a poor XY-correlation (r=0.6) between *Satb1* transcript and SATB1 protein levels **(Figure S2.F)**. Further analysis of our SATB1 immunoblots in late LPS-activated cells revealed at least two SATB1 bands **(Figure 2.F)**, suggesting either a loss of post-translational modification or the presence of multiple SATB1 isoforms. We estimated their apparent molecular weight using an ImageJ script (Ohgane, 2024). These bands correspond to apparent molecular weights of approximately 103 kDa (top) and 96 kDa (bottom). The unexpected presence of two bands prompted us to investigate SATB1 isoform expression in B-cells.

**Figure 2.**
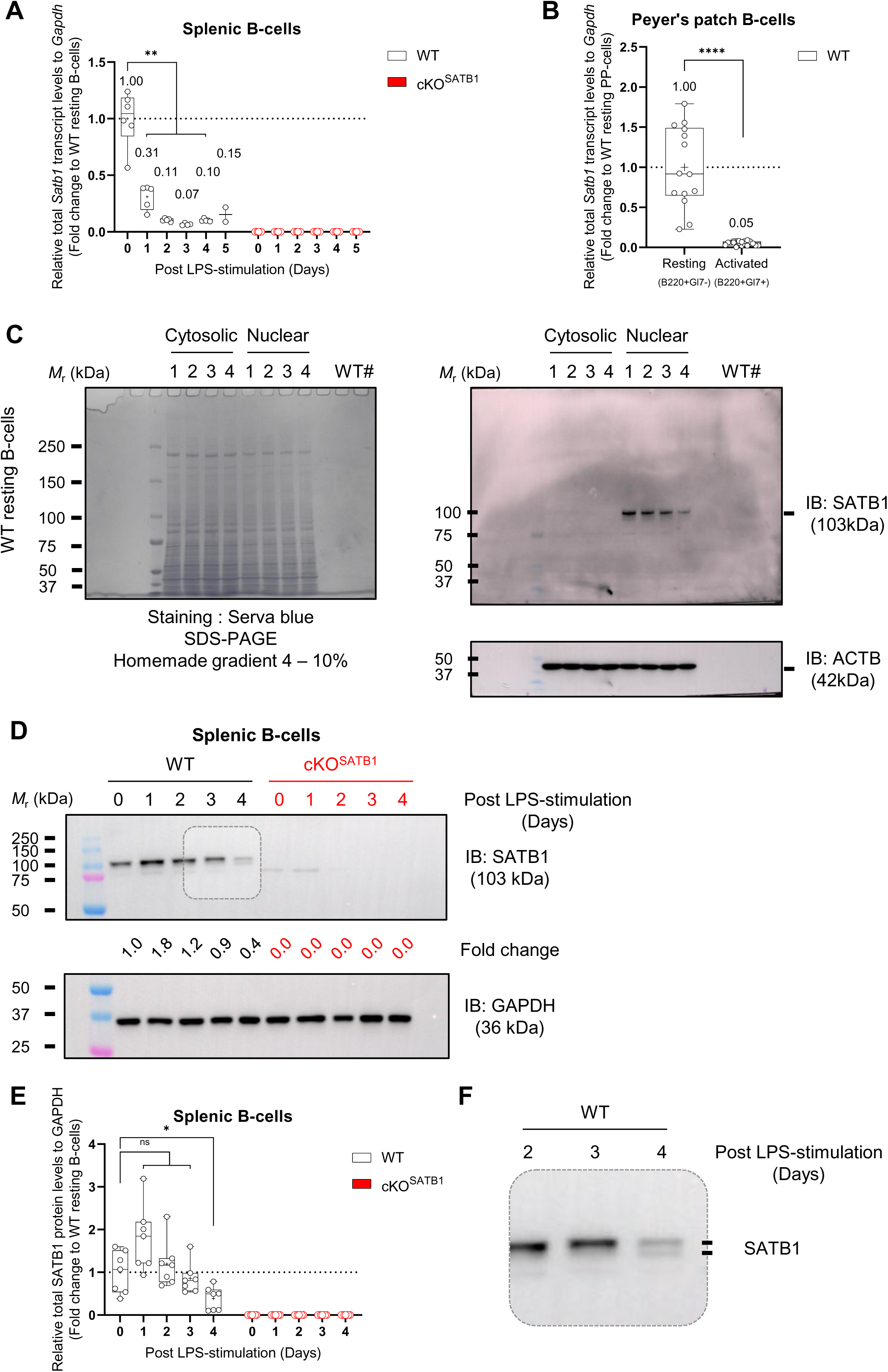
Divergent kinetics of *Satb1* transcript and SATB1 protein levels in mature B-cells. **(A)** Whisker plot showing quantitative RT-PCR analysis of total *Satb1* expression using primers shown in Figure S2.A. Expression was normalized to *Gapdh* and fold change was calculated relative to splenic resting B-cells. *n* = 6 for resting and LPS-activated days 1–4; *n* = 2 for LPS-activated day 5. **(B)** Whisker plot represents quantitative RT-PCR analysis of total *Satb1* expression in Peyer’s patch B-cells. Normalization and fold change were performed as in *(A)*, with resting Peyer’s patch B-cells as the reference. n = 14 for resting (B220⁺/GL7-); *n* = 18 for activated (B220⁺/GL7+). **(C)** Sorted WT resting B-cells were fractionated into cytosolic and nuclear fractions and were resolved by SDS-PAGE (homemade 4–10% gradient) and stained with Serva Blue dye. Parallel immunoblots were probed with anti-SATB1 antibody (1:5,000, ab109122, Abcam), and anti-β-actin (1:5,000, #5125S, Cell Signaling Technology) was used as a loading control. Four independent WT biological replicates are shown. **(D)** Representative immunoblot of total SATB1 protein levels in WT and cKOSATB1resting and LPS-activated (up to 4 days after LPS stimulation). Immunoblotting was performed with anti-SATB1 antibody (1:5,000, ab109122, Abcam) and anti-β-actin (1:5,000, #5125S, Cell Signaling Technology) as a loading control.Total SATB1 protein levels were normalized to GAPDH, and fold change was calculated relative to WT resting B-cells (day 0). **(E)** Whisker plot showing relative quantification of SATB1 protein levels from the immunoblot analysis shown in *(D)*. *n* = 7. **(G)** Magnified view of the immunoblot from *(D)*. P-value was determined using a two-tailed Mann–Whitney U test, significant differences from WT only are indicated: P > 0.05 (ns); P < 0.05 (*); P < 0.01 (**).

### Full-length and alternatively spliced *Satb1* isoforms are expressed in mature B-cells

We further analyzed our RNA-seq data (Thomas et al., 2023) and plotted them in sashimi plots to visualize exon-exon junction reads in resting and *in vitro* activated B-cells. We detected exon 11 skipping in resting and LPS-activated B-cells, similar to previously described in T-cells (Zelenka et al., 2023). In addition, we also identified exon 9 skipping specifically in resting B-cells **(Figure 3.A)**. To confirm the results obtained from sashimi plots, we used a reverse primer in exon 12 and a forward primer in exon 7 to amplify any potential transcript from this region. Among the four isoforms, we readily identified three of them at the expected sizes (f*ull*-*length*, *Δ11*, and *Δ9Δ11)* in two different mouse strains **(Figure 3.B)**. We also used a similar strategy with a different forward primer in exon 8 and found the same three isoforms at their expected sizes **(Figure 3.C)**. To approximate the relative quantity of each *Satb1* isoform, we measured band intensities of amplified *Satb1* isoforms from both RT-PCR strategies. In both approaches, the *Satb1 Δ11* isoform represented the main transcript, whereas the *full-length* and *Δ9Δ11* were minorly expressed **(Figure S3.A)**. Next, to specifically quantify each *Satb1* isoform, we designed forward and reverse primers that overlap the exon of interest **(Figure S3.B, left panel)**. Since there was a significant risk that a single primer could anneal the single exon 10 and amplify from either cDNA or genomic DNA (gDNA), we verified that RT-PCR performed on B-cell cDNA produced a single specific band, whereas using gDNA as a template lead to multiple unspecific products **(Figure S3.B, top right panel)**. Furthermore, melting curves analysis confirmed the reliability of our RT-qPCR reactions **(Figure S3.B, bottom right panel)**. Based on these validated assays, we quantified all *Satb1* splicing isoforms in resting and activated B-cells, both *in vivo* and *in vitro*. All *Satb1* splicing isoforms were detected and were downregulated upon LPS activation and in activated Peyer’s patch B-cells **(Figure 3.D)**, consistent with the overall *Satb1* downregulation observed in our previous RT-qPCR data **(Figures 1-2)**. Similar results were obtained when normalizing to *Actb* **(Figure S3.C)**. Analysis of publicly available data from NCBI Genome Data Viewer (RefSeq GCF_000001635.27-RS_2024_02) revealed one validated *Satb1* transcript that harbors skipping of exon 9 alone (NP_001344569.1). This unexpected finding expanded the possible outcomes of *Satb1* alternative splicing to four mRNA isoforms: *Full* (exon 11 retained), *Δ11* (exon 11 skipped; referred to as the short isoform in Zelenka et al., 2023), *Δ9* (exon 9 skipped, exon 11 retained), and *Δ9Δ11* (both exons 9 and 11 skipped) as detailed in **Figure 4A**. Interestingly, the two novel transcripts (*Δ9* and *Δ9Δ11*) maintain an open reading frame and could potentially give rise to functional SATB1 proteins with different domain conformation (**Figure 4B**). Next, we aimed to identify each SATB1 isoform in B-cells, for that we employed an immunoprecipitation (IP) approach using either an antibody targeting the N-terminal region of SATB1, a domain conserved across all known and putative isoforms or an antibody specific to the full-length SATB1 isoform **(Figure 5.A)**, raised against the exon 11-encoded extra peptide and previously validated (Zelenka et al., 2023). We compared this full-length-specific antibody to the pan-SATB1 antibody for IP from nuclear extracts of resting and *in vitro* LPS-activated WT and cKO^SATB1^ B-cells, as well as from whole-cell lysates of WT thymus (positive control). Following IP, input (fraction #1), MOCK (fraction #2), and eluted fractions (fraction #3) were resolved by SDS-PAGE and immunoblotted (IB) with the an antibody that reveal all SATB1 isoforms **(Figure 5.A in grey)**. In WT samples, SATB1 was readily detected in the eluted fractions (#3) from both IP antibodies in resting **(Figures 5.B–S4.A)**, in LPS-activated B-cells **(Figures 5.C–S4.B)**, and thymus **(Figure 5.D–S4.C**). As expected, SATB1 was absent in cKO^SATB1^ controls (**Figures 5 and S5**). Collectively, these findings establish that the full-length SATB1 isoform is expressed not only in T-cells, but also in resting and *in vitro* activated B-cells.

**Figure 3.**
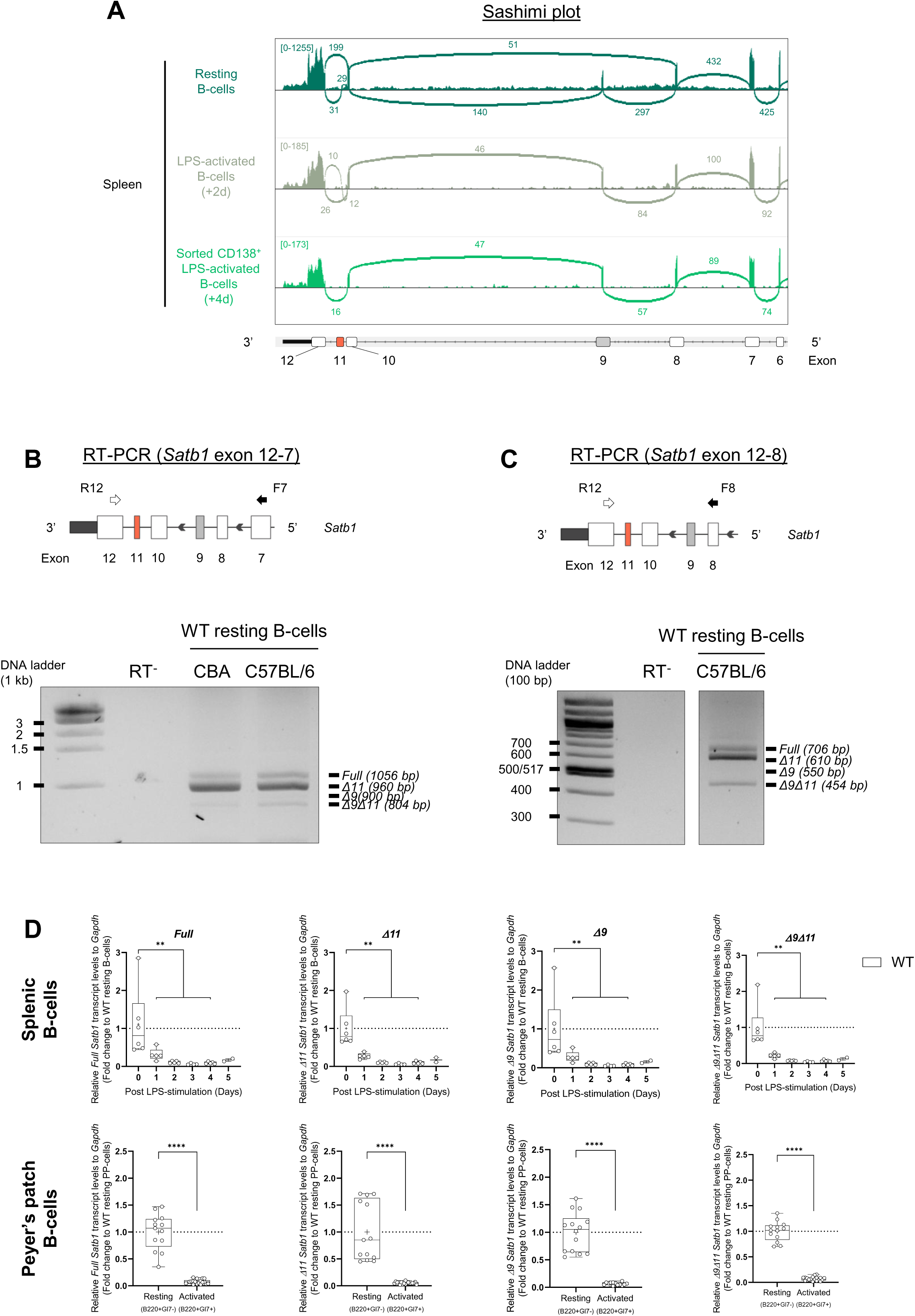
*Full-length* and alternatively spliced *Satb1* isoforms are expressed in mature B-cells. **(A)** Sashimi plots generated from RNA-seq data (BAM files) of resting B-cells, LPS-activated B-cells (day2) and in plasmablast CD138 positives cells, sorted after 4 days of LPS stimulation showing exon–exon junctions across the *Satb1* locus using IGV (v2.19.7). Read counts for are shown as mean values: *n* = 4 for resting and LPS-activated day-2 B-cells; *n* = 2 for plasmablast CD138 positives. **(B) Top panel :** schematic representation of primer annealing sites: a reverse primer in exon 12 (R12, white arrow) and a forward primer in exon 7 (F7, black arrow). **Bottom panel :** representative 1.5% agarose gel with separation of potential spliced isoform bands in WT resting splenic B-cells from two mouse genetic backgrounds (CBA and C57BL/6). **(C) Top panel :** schematic representation of primer location : a reverse primer in exon 12 (R12, white arrow) and a forward primer in exon 8 (F8, black arrow). Bottom panel : representative 2% agarose gel with the satb1 spliced isoforms **(D)** Whisker plot showing quantitative RT-PCR analysis of each *Satb1* isoform. Expression was normalized to *Gapdh* and fold change was calculated resting B-cells (**upper panel**) and to resting Peyer’s patch B-cells (**lower panel**). For splenic B-cells: = 6 for resting and LPS-activated days 1–4; *n* = 2 for LPS-activated day 5. For Peyer’s patch B-cells: *n* = 13 for resting (B220⁺/GL7⁻); *n* = 18 for activated (B220⁺/GL7+). P-value was determined using a two-tailed Mann–Whitney U test for only significant differences are indicated: P < 0.0001 (****).

**Figure 4.**
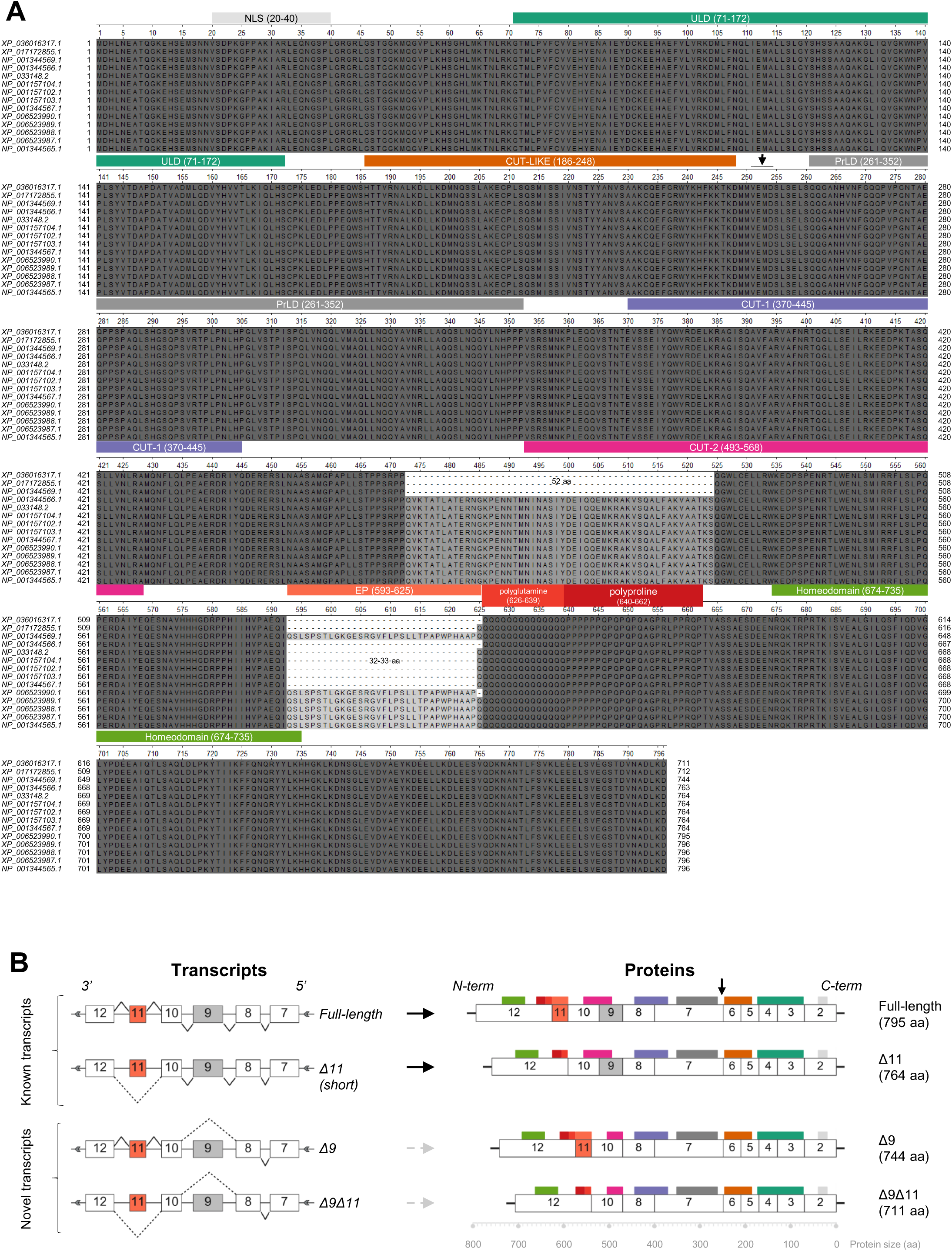
Protein structure of murine SATB1 proteins. **(A)** Alignment of 14 murine SATB1 protein sequences from NCBI (RefSeq GCF_000001635.27-RS_2024_02), including 8 validated (annotated: NP) and 6 experimental (annotated: XP) sequences, performed using Unipro UGENE software (v53.0). Exon 9 skipping appears as a gap of 52 amino acids, and exon 11 skipping as a gap of 32–33 amino acids. Protein domains are color-coded as follows: NLS (light gray), ULD (dark green), CUT-LIKE (brown), prion-like domain (dark gray), CUT-1 (purple), CUT-2 (pink), extra peptide; EP (light orange), polyproline; polyP (dark orange), polyglutamine; polyQ (dark red), homeodomain; HD (light green). The reported caspase-6 cleavage site is indicated at position 254 aa. Amino acid positions for each domain are shown next to the domain name. **(B)** Schematic representation of known (*full-length* and *short*, referred to as *Δ11*) and novel (*Δ9* and *Δ9Δ11*) *Satb1* splicing isoforms. For clarity and readability, only the N-terminal region (encoded by exons 7–12) of each transcript is shown, rather than the full-length sequence and the corresponding of known and putative SATB1 protein.

**Figure 5.**
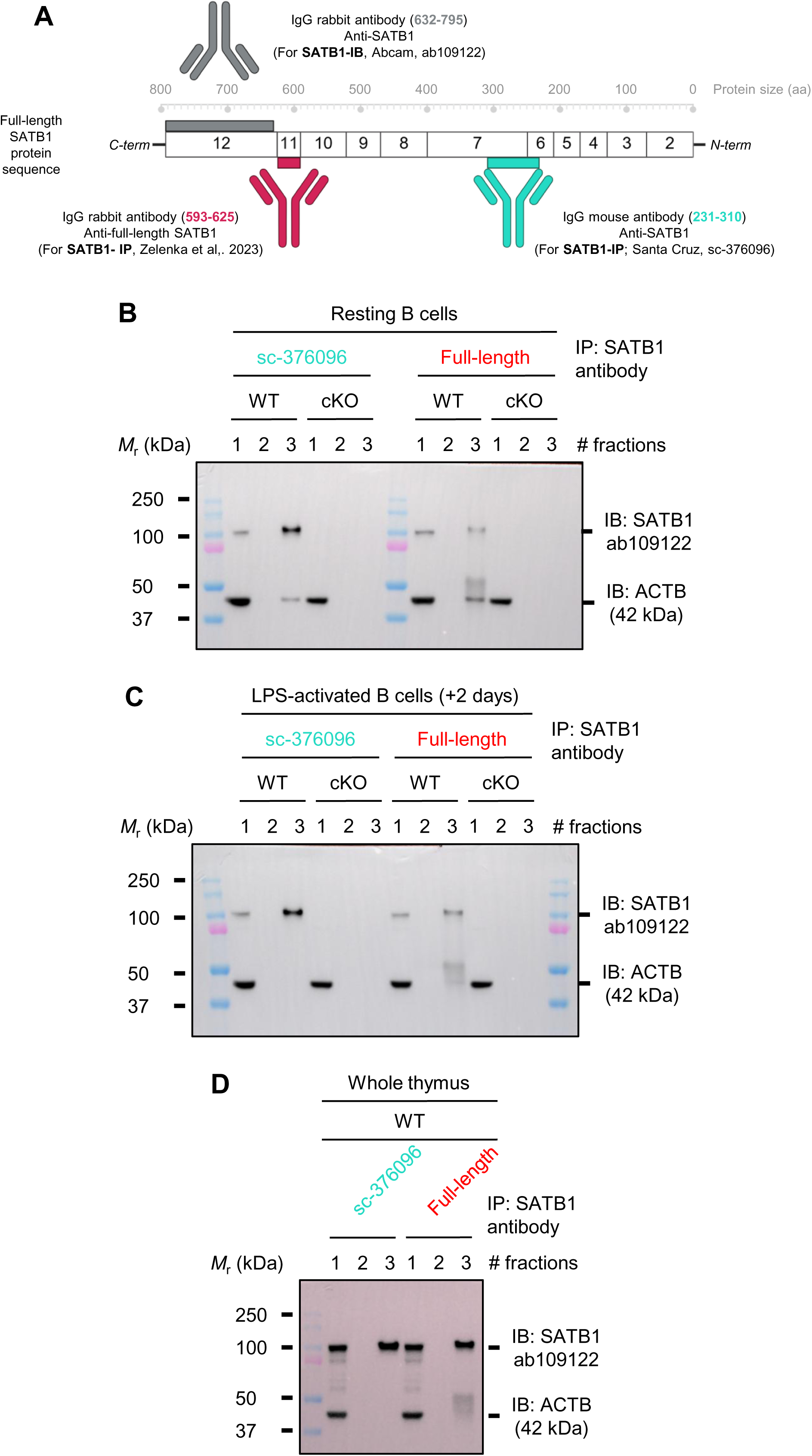
Full-length SATB1 is expressed in both lymphocyte lineage, B- and T-cells. **(A)** Schematic representation showing the binding site of three antibodies targeting SATB1 for immunoprecipitation (IP) and immunoblotting (IB) experiment. Two antibodies are used for SATB1 pull-down: the monoclonal mouse anti-SATB1 antibody (sc-376096, Santa Cruz), which targets the N-terminal region of fulllength SATB1 (indicated in red); and the polyclonal rabbit anti-full-length SATB1 antibody (indicated in cyan). The third antibody is used for SATB1 immunoblotting (ab109122, Abcam) indicated in grey. **(B-D)** Immunoprecipitation (IP) of SATB1 from nuclear extracts of resting **(B)** and LPS-activated **(C)** WT and cKOSATB1 splenic B-cells, and from whole-cell lysates of WT thymus **(D)**.

### SATB1 forms a high-molecular-weight protein complex exceeding 700 kDa in mature B-cells

We next asked whether SATB1 exists as a monomer (∼103 kDa) or as part of a larger native assembly. To address this, we assessed its migration through native polyacrylamide gels. We first resolved whole-cell lysates from resting B-cells by commercial blue native PAGE (BN-PAGE). In resting B-cells, SATB1 was detected above the 720 kDa molecular weight marker **(Figure S5.A)**, indicating the presence of a large native complex. To exclude the possibility that this low electrophoretic mobility was due to chromatin tethering (rather than genuine protein–protein interactions), we treated nuclear extracts with benzonase to digest nucleic acids. Benzonase treatment did not drastically alter the migration of the native SATB1 complex **(Figure S5.B)**, ruling out DNA- or RNA-mediated retention. We confirmed these findings using homemade 4–16% native PAGE with benzonase-treated nuclear extracts: SATB1 was again detected as a high-molecular-weight complex in WT resting B-cells, but absent in cKO^SATB1^ controls **(Figure 6.A)**. To further verify that this native complex corresponds to a bona fide SATB1-containing complex rather than a non-specific aggregate, we performed two-dimensional electrophoresis (native PAGE followed by denaturing SDS-PAGE). The SATB1 signal resolved as a single spot migrating from the native gel into the denaturing gel (white arrows in **Figure 6.B** and **Figure S6.C**). As expected, this complex was absent in cKO^SATB1^ controls **(Figure 6.B)**. These results confirmed that SATB1 exists as a stable, high-molecular-weight protein complex (>720 kDa) in resting B-cells.

**Figure 6.**
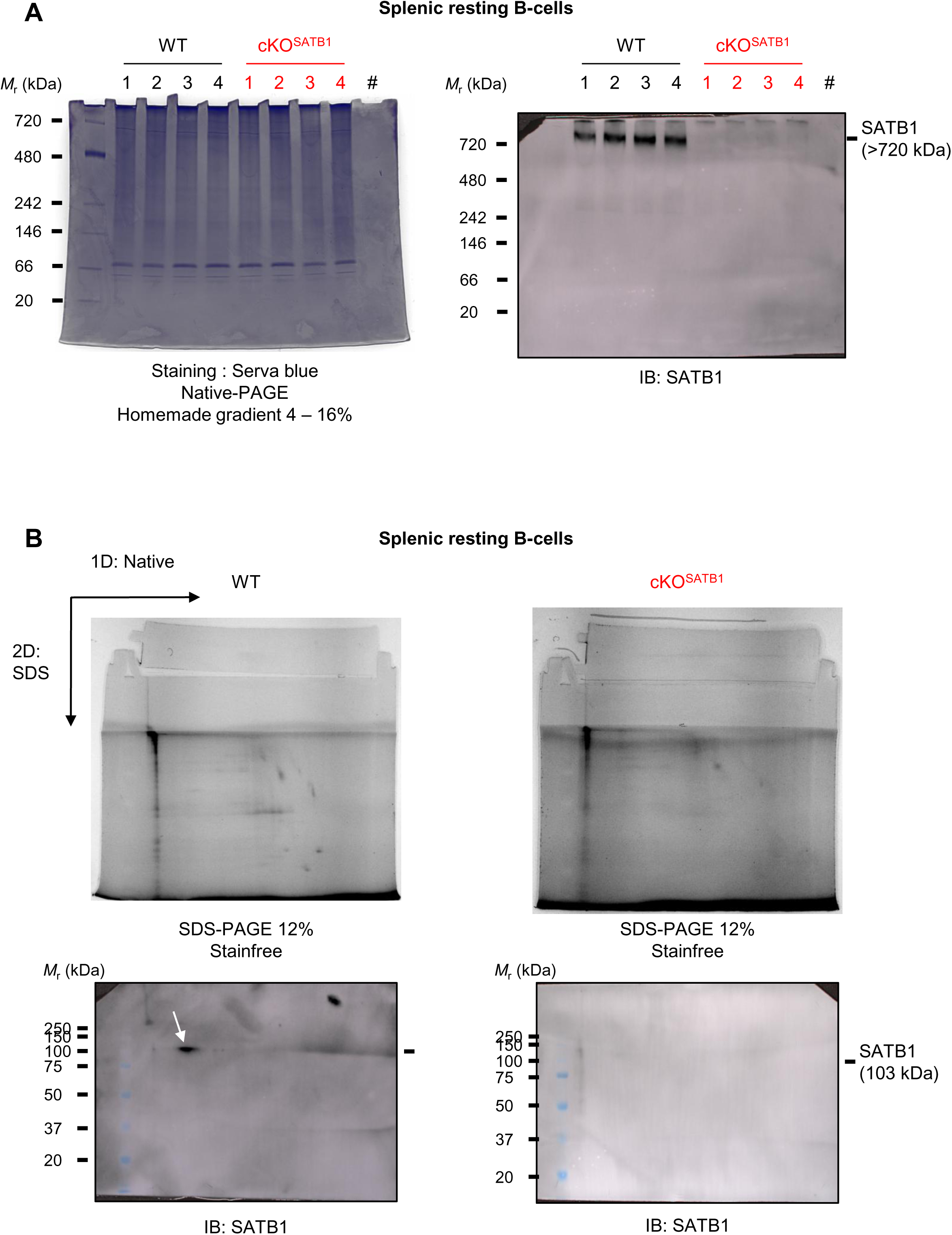
SATB1 assembles into a native >720 kDa complex in mature B-cells. **(A)** Nuclear extracts from WT and cKOSATB1 resting B-cells were resolved by native PAGE (homemade 4–16% gradient). One gel was stained with Serva Blue to visualize total protein, while a parallel gel was immunoblotted with anti-SATB1 antibody (1:5,000, ab109122, Abcam). **(B)** Benzonase-treated nuclear extracts (10 μg) from WT resting B cells were first resolved by native PAGE (4–16% gradient; first dimension). The entire native lane was excised and subjected to denaturing SDS-PAGE (12%; second dimension). Total protein was visualized using stain-free technology (Bio-Rad; top panel, followed by immunoblotting with anti-SATB1 antibody (bottom panel), revealing a single SATB1 spot (white arrow).

### RNA-binding proteins are the predominant SATB1 interactors in resting and activated B-cells

Given that SATB1 is a high-molecular-weight protein complex in B-cells, we next sought to characterize its interacting partners. For this purpose, we employed an immunoprecipitation (IP) approach using an antibody targeting the N-terminal region of SATB1. Previously, we established that the Santa Cruz antibody captures both known SATB1 isoforms present in the nucleus and that the Abcam detection-antibody reliably recognized the full-length isoform **(Figure 5)**. Thus, we used these suitable antibodies in our co-IP/MS approach. The complete IP workflow is detailed in **Figure S6.** Nuclear extracts were pooled and first incubated with Dynabeads® protein A alone (MOCK), followed by incubation with SATB1 antibody-conjugated beads for IP. All fractions were collected to monitor SATB1 elution by immunoblotting with the anti-SATB1 antibody (ab109122, Abcam). SATB1 was clearly detected in fractions 7 and 9, but not in fraction 2 **(Figures 7.A and S7A)**, confirming that most of the target protein was efficiently bound to the antibody. Western blotting validated the presence of SATB1 in WT IP eluates **(Figure S7.A, right panel)** and its absence in cKO^SATB1^ controls **(Figure S7.B, right panel)**.

**Figure 7.**
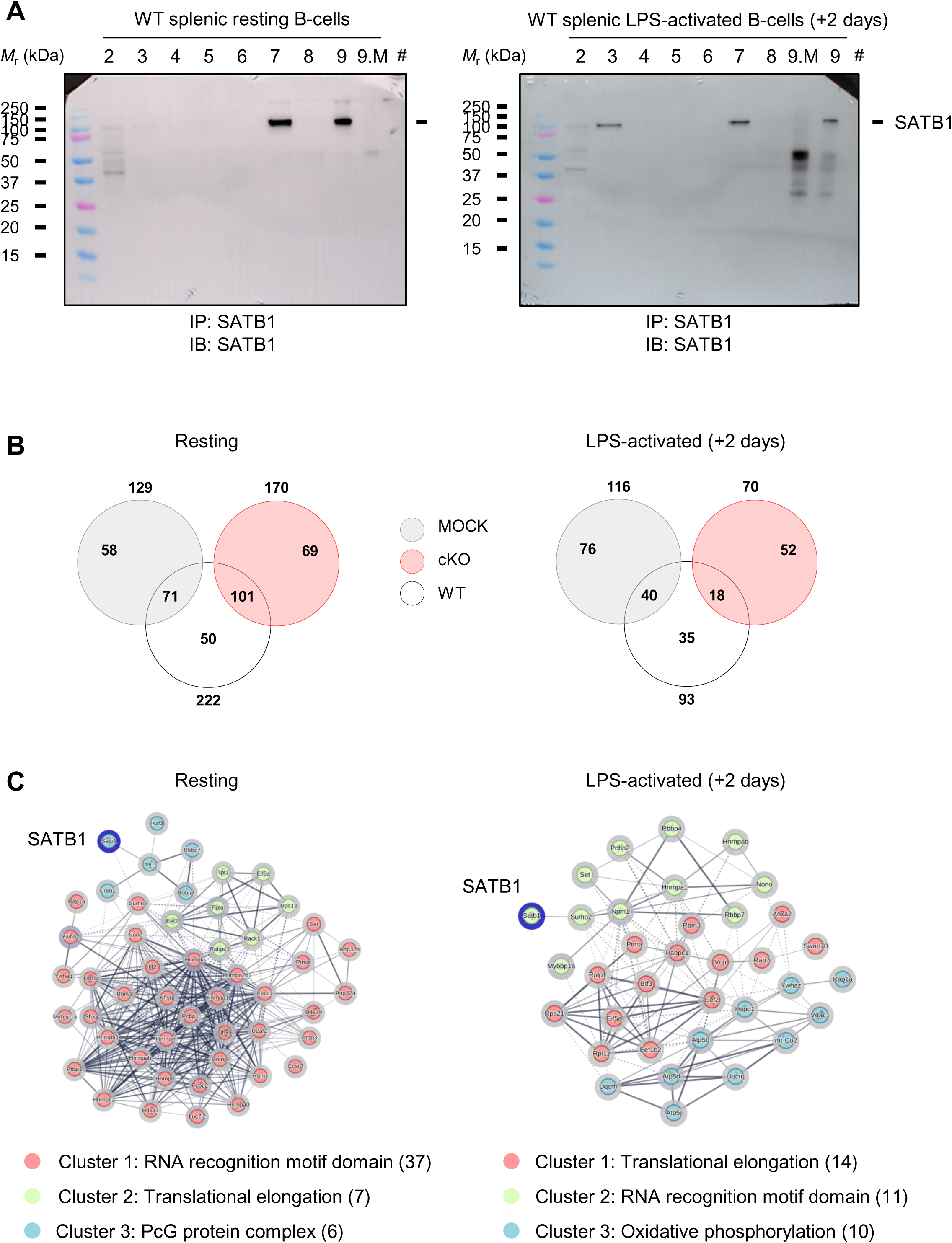
RNA-binding proteins are the predominant SATB1 partners in resting and LPS-activated B-cells. **(A)** Western blot analysis of SATB1 (1:5,000, ab109122, Abcam) in collected fractions from the IP experiment using WT splenic B-cells (resting and day-2 LPS-activated). Fraction numbers (#) correspond to those described in the Material and Methods **(B)** Venn diagram showing the average distribution of SATB1 interactors found in MOCK (#Fraction 7.M), cKO and WT (#Fraction 7) in resting and LPS-activated B-cells (+ 2 days). *N* = 3 for WT; *N* = 2 for cKOSATB1. **(C)** Clustering analysis of SATB1-interacting proteins identified by mass spectrometry in resting and LPS-activated B-cells (STRING, k-means clustering, *N* = 3).

The MOCK IP (fraction 7.M) and SATB1 IP eluates (fraction 7) from resting and in vitro LPS-activated WT and cKO^SATB1^ B-cells were subjected to mass spectrometry (MS) to characterize the SATB1 interactome. As expected, our IP approach was specific to SATB1 as we did not detect SATB2 peptides in our MS analysis. In resting B-cells, we identified a substantial number of SATB1-associated proteins, whereas in LPS-activated B-cells, the global numbers of interactors decreased. Strikingly, SATB1-associated proteins were predominantly enriched in RNA-binding factors in resting and *in vitro* activated B-cells **(Figure 7.B)**. These unexpected findings unveil a potential function for SATB1 in RNA regulation that warrants further investigation in B-cells.

## Discussion

In this study, we uncovered multilayered regulation of SATB1 in murine B-cells, revealing that *Satb1* expression is controlled through alternative promoter usage and, alternative splicing. Our findings establish that *Satb1* promoters are differentially regulated in B-cells compared to T-cells. Among the four alternative promoters, P2 remains inaccessible and transcriptionally silent in B-cells whereas P1 and P3 undergo a dynamic switch upon LPS-induced activation, shifting from P3 to P1. This lineage-specific promoter usage echoes the shift from P1 to P2 and P3 observed T-lineage cells upon *in vitro* activation (Khare et al., 2019; Patta et al., 2020),

The *Satb1* P4 promoter showed a forward transcription activity. Intriguingly, P4 remained the main site where transcription on the reverse strand occurs in B and T lymphocytes. Thus, P4 displays pervasive transcription on both strands, reminiscent of bidirectional promoter activity (Xu et al., 2009). This pervasive transcription activity could be dedicated to non-coding RNA transcription (Jensen et al., 2013; Poliseno et al., 2024), more specifically to an upstream region identified as the *Satb1* antisense transcript (*SATB1*-*AS1*), which might likely act as an enhancer RNA (eRNA) as described in humans (Wang et al., 2025). More importantly, the P4 promoter region matches a recently characterized ultra-conserved silencer region (Bai et al., 2025; Hansen and Hodges, 2022), suggesting an additional repressive function for *SATB1-AS1*. In contrast to humans, the putative function of such a *Satb1* antisens transcript has not yet been studied in *Mus musculus.* One tempting hypothesis could be that eRNAs contribute to chromatin organization, since involved in the nucleation and maintenance of nuclear condensates, helping to concentrate necessary components near regulatory elements such as super-enhancers (Arnold et al., 2019; Statello et al., 2021; Wang et al., 2026).

A striking observation was the poor correlation between *Satb1* transcript and SATB1 protein levels. While *Satb1* transcripts rapidly declined following B-cell activation, protein levels transiently increased before gradually decreasing. The significant drop in total *Satb1* transcript suggests tight post-transcriptional regulation upon B-cell activation. The enhanced SATB1 protein stability remains enigmatic. While our study did not focused on SATB1 protein turnover, our data suggest that its stabilization is unlikely to depend on the ubiquitin-proteasome system (Patta et al., 2020; Tan et al., 2008), nor on cleavage by caspase-6 or caspase-3 as previously described (Galande et al., 2001; Notani et al., 2010; Olson et al., 2003; Sun et al., 2006; Tan et al., 2008). Indeed, we did not observe in our immunoblots any cleavage fragments as the consequence of SATB1 degradation. We were also unable to detect in our co-IP/MS analysis any factor, caspases or PML proteins, linked to cleavage or nuclear relocation. We hypothesize that the discrepancy between transcription and protein levels is more likely to be the consequence of a tight *Satb1* transcripts translation efficiency. Indeed, 3′UTR lengths of *Satb1* transcripts were proposed to modulate translation efficiency in T cells as suggested by Patta et al. (2020). The relative stability of SATB1 protein in late-activated B cells raises the question of its potential function at this stage, when chromatin remodelling and transcriptional reprogramming are critical.

Beyond the well-folded domains such as NLSn, CUT, CUT-Like, Ubiquitin-like and Homeodomain, SATB1 protein harbours several intrinsically disordered regions, including the extra peptide and low-complexity domains such as polyproline and polyglutamine stretches. This specific domain architecture allows the SATB1 homeodomain to recognize AT-rich sequences, while the CUT domains bind within the nucleosomal core with positive cooperativity (Ghosh et al., 2019), preferentially targeting nucleosome-dense regions—a feature that supports its candidacy as a pioneer transcription factor (Ghosh et al., 2019; Kohwi et al., 2025; Yasui et al., 2002; Zelenka et al., 2022). The intrinsically disordered nature of SATB1 suggests that its protein conformation is flexible, and thus its functions can drastically change. Decades ago, several groups showed two SATB1 bands in cell lines (Jurkat, K562, HEK293T) or thymocytes (de Belle et al., 1998; Dickinson and Kohwi-Shigematsu, 1995; Kaur et al., 2016; Nakagomi et al., 1994). However, one study in particular failed to detect *Satb1* spliced isoforms and ruled out this possibility (Cunningham et al., 1994), instead suggesting that the different masses resulted from post-translational modifications such as phosphorylation. While described as phosphorylated in T cells, no phosphorylated SATB1 (S635 for Δ11, S666 for full-length SATB1) was detected through our co-IP/MS in either resting or LPS-activated B-cells (Notani et al., 2010; Pavan Kumar et al., 2006; Tan et al., 2010; Zelenka et al., 2023). The lower molecular weight SATB1 band appearing in LPS-activated B-cells suggests either a loss of post-translational modification and/or the expression of a shorter protein isoform. Given the absence of phosphorylated SATB1 in resting B cells, the lowed band cannot be explained by SATB1 dephosphorylation. Although our co-IP/M experiment identified SUMO2 as an interactor of SATB1 in resting B and activated B cells; it cannot be excluded that SUMOylated SATB1 corresponds to the upper band and that the lower band is the consequence of the loss of this post-translational modification.

More recently, a study showed that two (short and long) SATB1 isoforms are expressed in thymus (Zelenka et al., 2023). Instead, we detected four *Satb1* splicing isoforms in B-cells, including two novel transcripts (*Δ9* and *Δ9Δ11*). Interestingly, while the *Δ9* isoform (NP_001344569.1) is a validated splicing isoform in the NCBI database expressed in T cells and neurons, we failed to detect consistent amounts of this variant transcript in splenic B cells, suggesting a very low expression.

AlphaFold predictions of the already described full-length and Δ11 protein isoforms in B and T cells, as well as the newly identified *Δ9Δ11* and putative *Δ9* variant transcripts showing distinct conformations with specific biophysical properties (**Figure S8**). Interestingly, protein isoforms devoid of exon 9, removes part of the CUT-2 domain and altered DNA- or protein-protein interaction. One study in particular provided evidence that the CUT-2 domain and the Q15 stretch are dispensable for DNA-binding activity (Nakagomi et al., 1994). However, SATB1 belongs to a family of proteins that harbour two CUT domains, such as CUX1 and CUX2 in mammals (Ramdzan et al., 2021). Interestingly, such proteins are often involved in DNA repair, since both CUT1 and CUT2 domains promote interaction with accessory base excision repair factors (e.g., OGG1 DNA glycosylase), but not through the homeodomain (Kaur et al., 2016), suggesting a potential function linking DNA-binding activity to DNA repair. The involvement of SATB1 in DNA repair remains an intriguing question in B cells, especially since our group recently proposed SATB1 as a negative regulator of somatic hypermutation of immunoglobulin genes, a mechanism that requires error-prone repair (Martin et al., 2023; Thomas et al., 2023).

Unexpectedly, we observed in resting B-cells, SATB1 forms a high-molecular weight complex (above 720 kDa) under native conditions. This high-order assembly in native condition has not been previously described and highlights a novel structural feature of SATB1 in B-cells. Notably, the presence of a single SATB1 isoform within this complex suggests that SATB1 may assemble into a multimeric structure that could serve as a platform for recruiting its interacting partners. In T cells, SATB1 recruit chromatin-modifying enzymes such as histone deacetylase 1 (HDAC1) to mediate transcriptional repression (Kumar et al., 2005), or associate with p300/CBP to promote gene activation (Fujii et al., 2003; Wen et al., 2005). However, our co-IP/MS analysis revealed that SATB1 associates predominantly with RNA-binding factors in B-cells, rather than mainly with chromatin-remodelling complexes (Fujii et al., 2003; Notani et al., 2010; Pavan Kumar et al., 2006; Purbey et al., 2009; Yasui et al., 2002; Zelenka et al., 2023). To note, some SATB1 protein partners recently identified in T cells also belong to the mRNA splicing pathway (Zelenka et al., 2023). Such a function in B and T cells in RNA splicing is consistent with its prion-like domains and its ability to form high-molecular-weight complexes. In the past, many groups failed to observe homodimerization of SATB1 (Nakagomi et al., 1994; Purbey et al., 2008; Tan et al., 2010; Yamaguchi et al., 2006; Yamasaki et al., 2007), although our data suggest that resting B cells contain a native complex composed of a single SATB1 isoform. Benzonase treatment suggests that SATB1 complex is stable independently of DNA- or RNA-binding activity. However, the precise stoichiometry and composition of the >720 kDa complex remain to be determined. Future studies employing super-resolution imaging will be essential to define the spatial organization of SATB1 with RNA-binding proteins, such as splicing factors. Given the proposed function for SATB1 in T cells, it is mandatory to further explore if SATB1 may also facilitate liquid-liquid phase separation in B cells.

In conclusion, we propose that SATB1 functions as a nexus between chromatin organization and RNA-associated functions in B-cells, with its activity tuned by alternative promoters, splicing, and post-transcriptional regulation. This dual functionality may explain the phenotypic differences between SATB1 functions in T and B lymphocytes, and underscores the adaptability of this pleiotropic protein to lineage-specific demands. SATB1 exemplifies how a single transcription factor can integrate multiple molecular functions—DNA- and RNA-binding, chromatin looping, nuclear matrix tethering, and recruitment of enzymatic activities—to orchestrate complex developmental programs. SATB1’s ability to function as both an activator and repressor, to bind both euchromatic and nucleosome-dense regions, to respond to post-translational modifications, and to act as both a DNA- and RNA-binding protein, reflects remarkable functional plasticity—a requirement in chromatin organization and likely in RNA-associated regulation, importantly in B-cells.

## Acknowledgments

We thank BISCEm and the animal core facility team for help with mouse work on both practical and regulatory aspects. JYF was supported by PhD fellowships from CNRS. LM was supported by la Société Française des Microscopies.

This work was supported by Fondation pour la Recherche Médicale, CNRS International Emerging Actions 2023, Partenariat Hubert Curien France/Grèce 2023, La Ligue Contre le Cancer (comités 87, 23).

## Author contribution

JYF, LM performed the experiments. TM manage the breeding of mice. EmP performed the MS experiments. TF and CS provided protocols and antibodies and participated to scientific discussion and gave their feedback on manuscript. EP and SLN designed and supervised the study JYF, EP and SLN wrote the manuscript.

---

**Figure S1.**
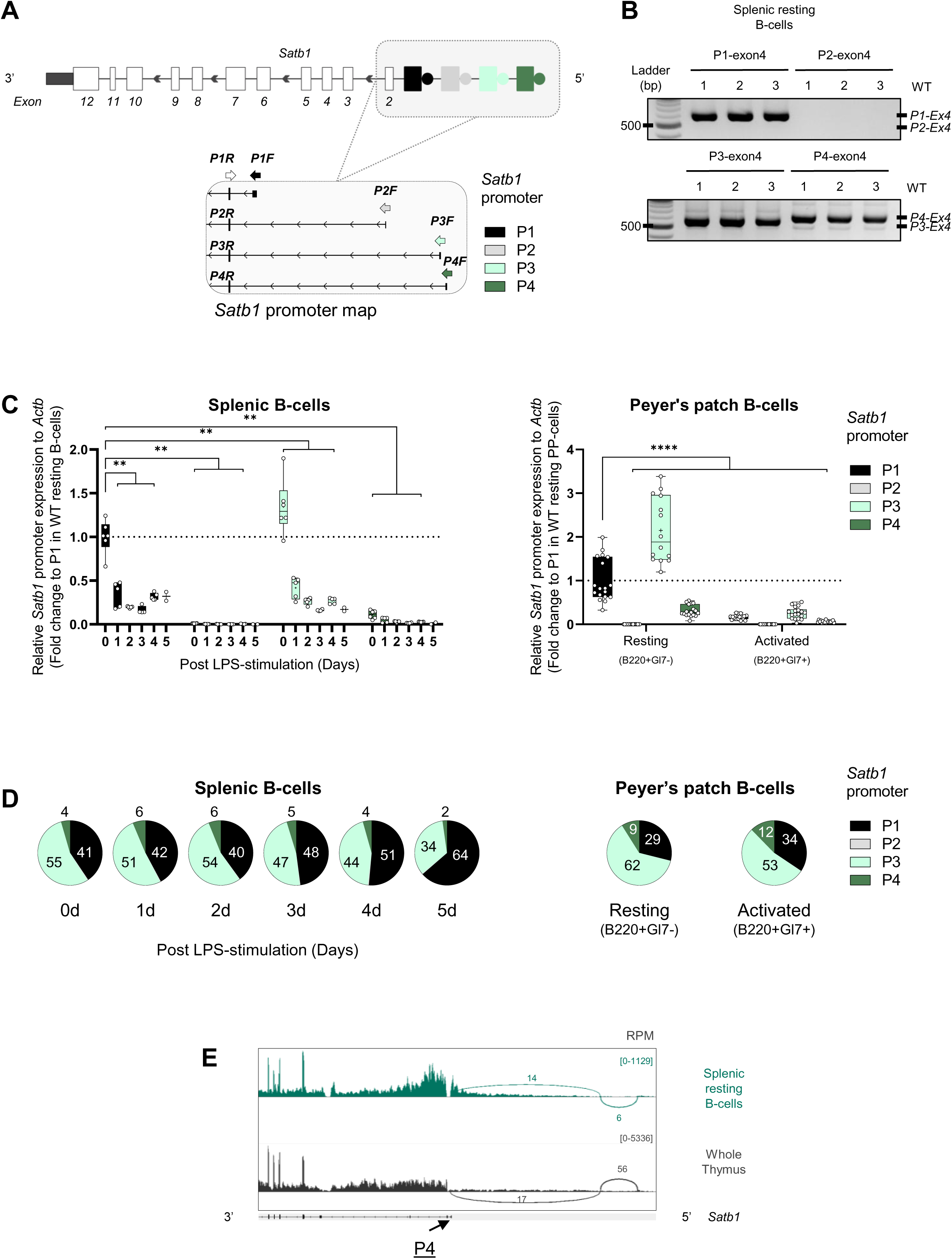
Dynamic of *Satb1* promoter activity in mature B-cells. **(A)** Schematic representation of the *Satb1* gene and primer locations used to quantify transcript levels from each *Satb1* promoter (Patta et al., 2020). **(B)** Agarose gel showing amplicons from RT-PCR to amplify forward *Satb1* cDNA from each alternative promoter to exon 4 in WT splenic resting B-cells. Three independent cDNA templates were used, and amplified products were resolved on a 3% agarose gel. **(C)** Whisker plots showing quantitative RT-PCR analysis of *Satb1* promoters in splenic **(right panel)** and Peyer’s patch **(left panel)** B-cells from WT mice similar to **Figure 1.C**. Expression was normalized to *Actb.* **(D)** Pie charts representing the quantitative RT-PCR data from **(C)** for each *Satb1* alternative promoter in splenic B-cells **(left panel)** and Peyer’s patch B-cells **(right panel)** relative to a total of 100%. **(E)** RNA-seq data showing junction on the reverse strand in splenic resting B-cells and whole thymus (GSE173470). Representative of *n* = 4 for resting B-cells and *n* = 3 for whole thymus. P-value was determined using a two-tailed Mann–Whitney U test; only significant differences are indicated: P < 0.01 (**); P < 0.0001 (****).

**Figure S2.**
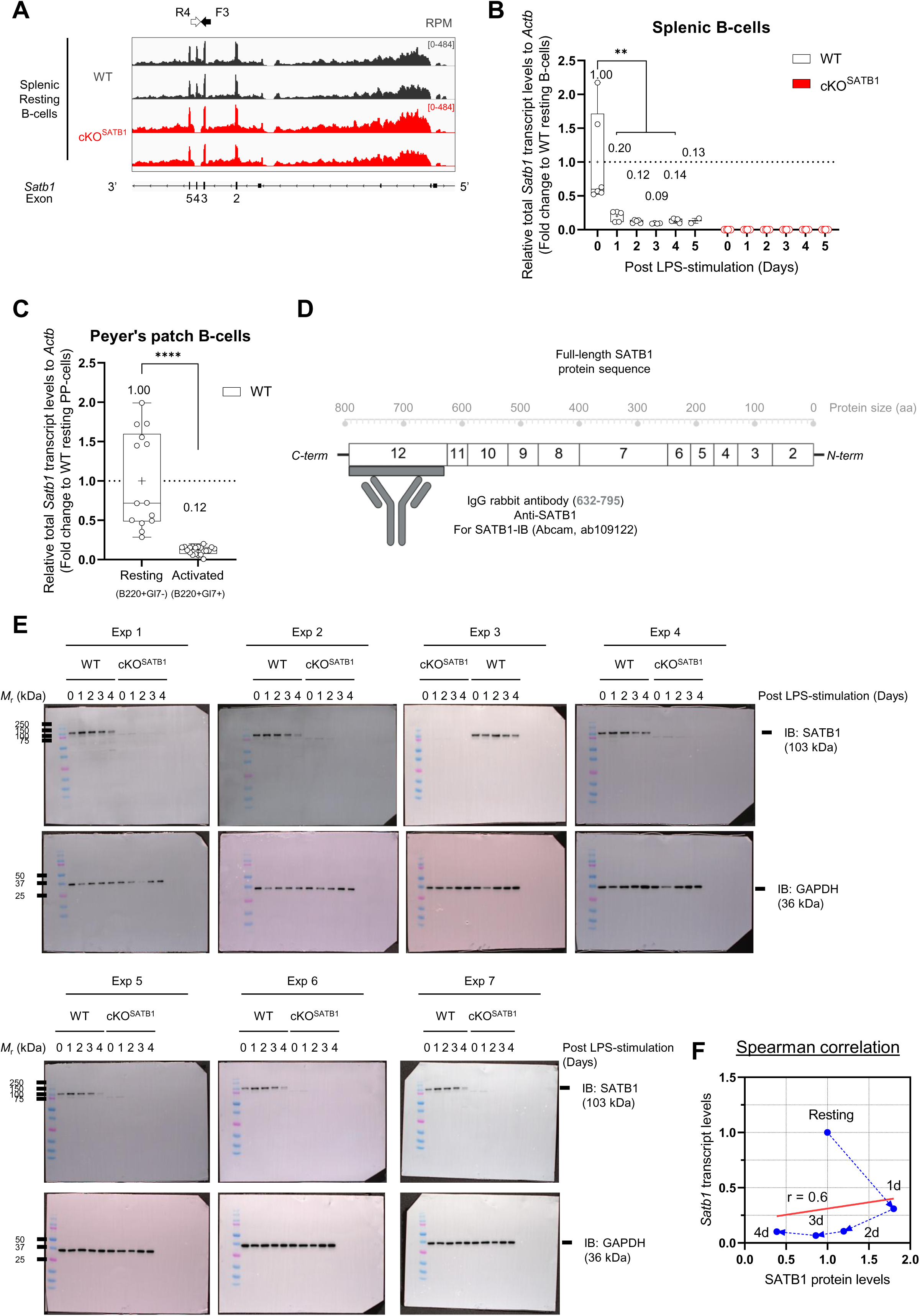
Divergent kinetics of *Satb1* transcript and SATB1 protein levels in mature B-cells. **(A)** Representative Bigwig from RNA-seq of WT (black) and cKOSATB1 (red) in resting B-cells. Arrows indicate the reverse primer in exon 4 (white) and the forward primer in exon 3 (black) used to quantify total *Satb1* transcript. **(B)** Whisker plot showing quantitative RT-PCR analysis of total *Satb1* expression in splenic B-cells. Expression was normalized to *Actb*, and fold change was calculated to resting B-cells. *n* = 6 for resting and LPS-activated days 1–4; *n* = 2 for LPS-activated day 5. **(C)** Whisker plot showing quantitative RT-PCR analysis of total *Satb1* expression in Peyer’s patch B-cells. Normalization and fold change calculation were performed as in *(B)*, with resting Peyer’s patch B-cells as the reference. *n* = 14 for resting (B220⁺/GL7-); *n* = 18 for activated (B220⁺/GL7+). **(D)** Schematic representation of full-length SATB1 protein (795 AA). The binding site of the monoclonal rabbit anti-SATB1 antibody (ab109122, Abcam) is indicated in light gray. **(E)** All raw Western blot images (*n* = 7) used to quantify total SATB1 and GAPDH protein levels in splenic B-cells upon LPS-induced activation, comparing WT versus cKOSATB1 mice. **(F)** XY correlation plot of *Satb1* transcript levels (Y-axis, data from **Figure 2.A**) versus SATB1 protein levels (X-axis, data from **Figure 2.E**) in splenic B-cells from resting to day-4 LPS-activated cells. Spearman rank correlation (red fitted line): rs[5] = 0.6, p > 0.05 (ns). P-value was determined using a two-tailed Mann–Whitney U test, significant differences from WT only are indicated: P < 0.01 (**); P < 0.0001 (****).

**Figure S3.**
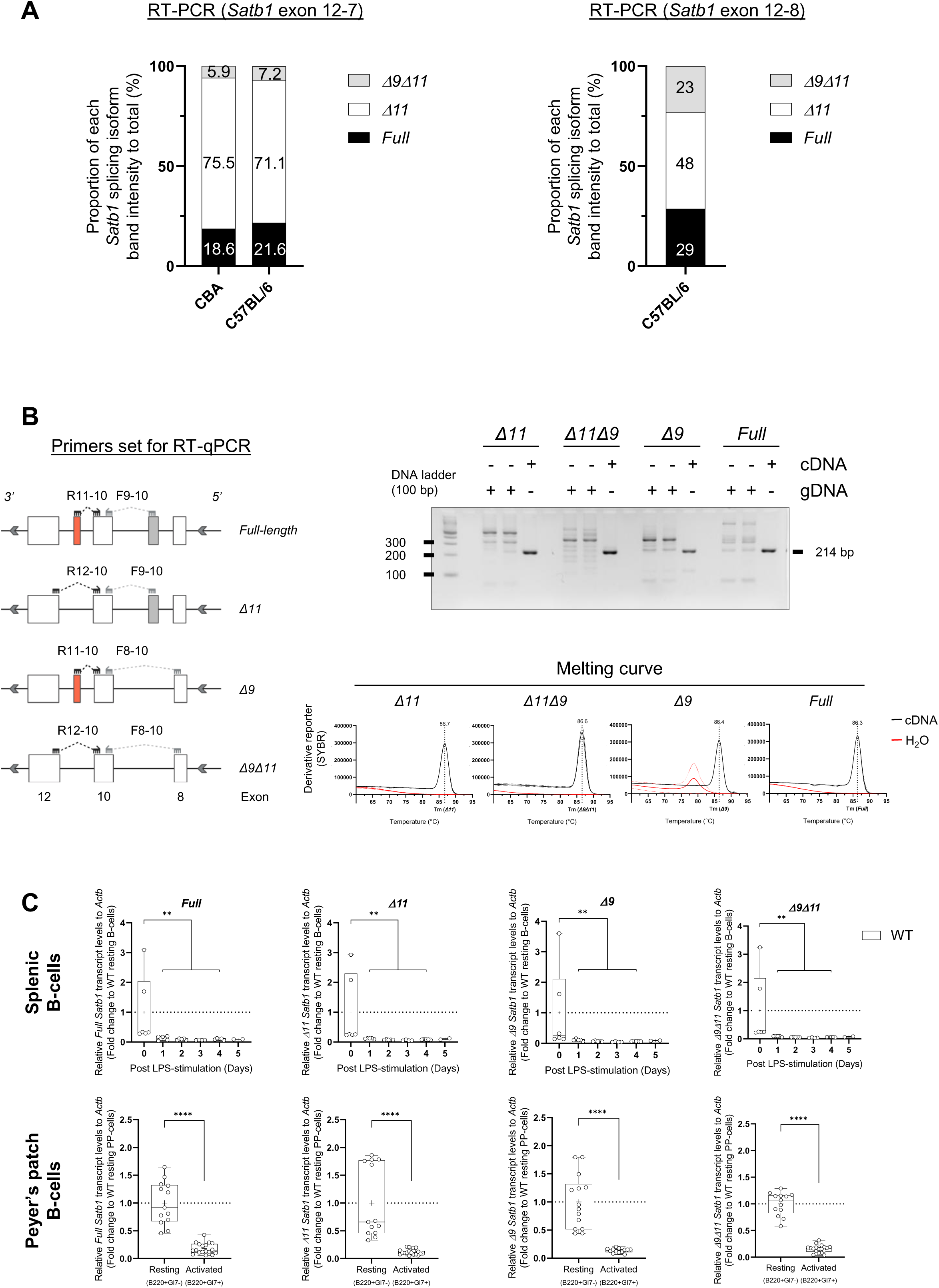
*Full-length* and alternatively spliced *Satb1* isoforms are expressed in mature B-cells. **(A) Left panel**: Bars graphs representing proportional band intensity of each *Satb1* splicing isoforms (*full-length*, *Δ11*, *Δ9* and *Δ9Δ11*) from Figure 3.B (total = 100%). Right panel : As in left from **Figure 3.C**. **(B) Left panel**: schematic representation of primers used to specifically detect each *Satb1* splicing isoform by quantitative RT-PCR and RT-PCR, using forward and reverse primers that overlap the exon of interest. **Right top panel**: Representative 3% agarose gel of RT-PCR performed with the designed primers using cDNA or gDNA as template. **Bottom right panel** : Representative melting curves of each q-RT-PCR strategies. **(C)** Whisker plot showing quantitative RT-PCR analysis of each *Satb1* splicing isoform. Expression was normalized to *Actb* and fold change was calculated to resting B-cells **(upper panel)** and to resting Peyer’s patch B-cells **(lower panel)**. For splenic B-cells: *n* = 6 for resting and LPS-activated days 1–4; *n* = 2 for LPS-activated day 5. For Peyer’s patch B-cells: *n* = 13 for resting; *n* = 18 for activated. P-value was determined using a two-tailed Mann–Whitney U test for only significant differences are indicated: P < 0.01 (**); P < 0.0001 (****).

**Figure S4.**
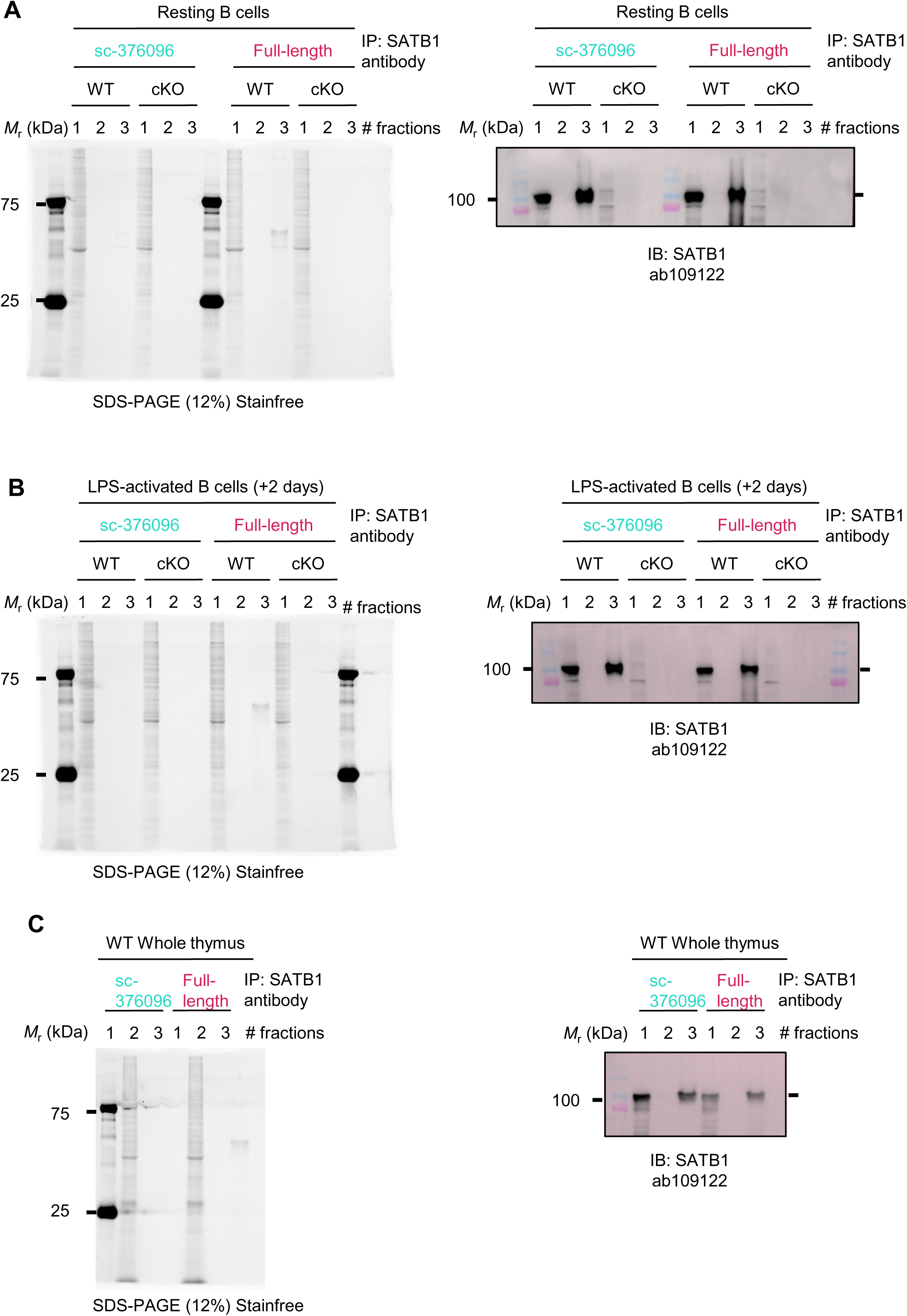
Full-length SATB1 is expressed in both B- and T-cells. **(A–C)** Immunoprecipitation (IP) fractions from WT and cKOSATB1 resting B-cells, LPS-activated B-cells (+ day 2), and WT thymus lysates were resolved by SDS-PAGE (12%). For each panel **(A–C)**: the left image shows total protein loading assessed by stain-free technology (Bio-Rad) prior to transfer; the right image shows the corresponding immunoblot of the same fractions after transfer, probed with anti-SATB1 antibody (1:5,000, ab109122, Abcam).

**Figure S5.**
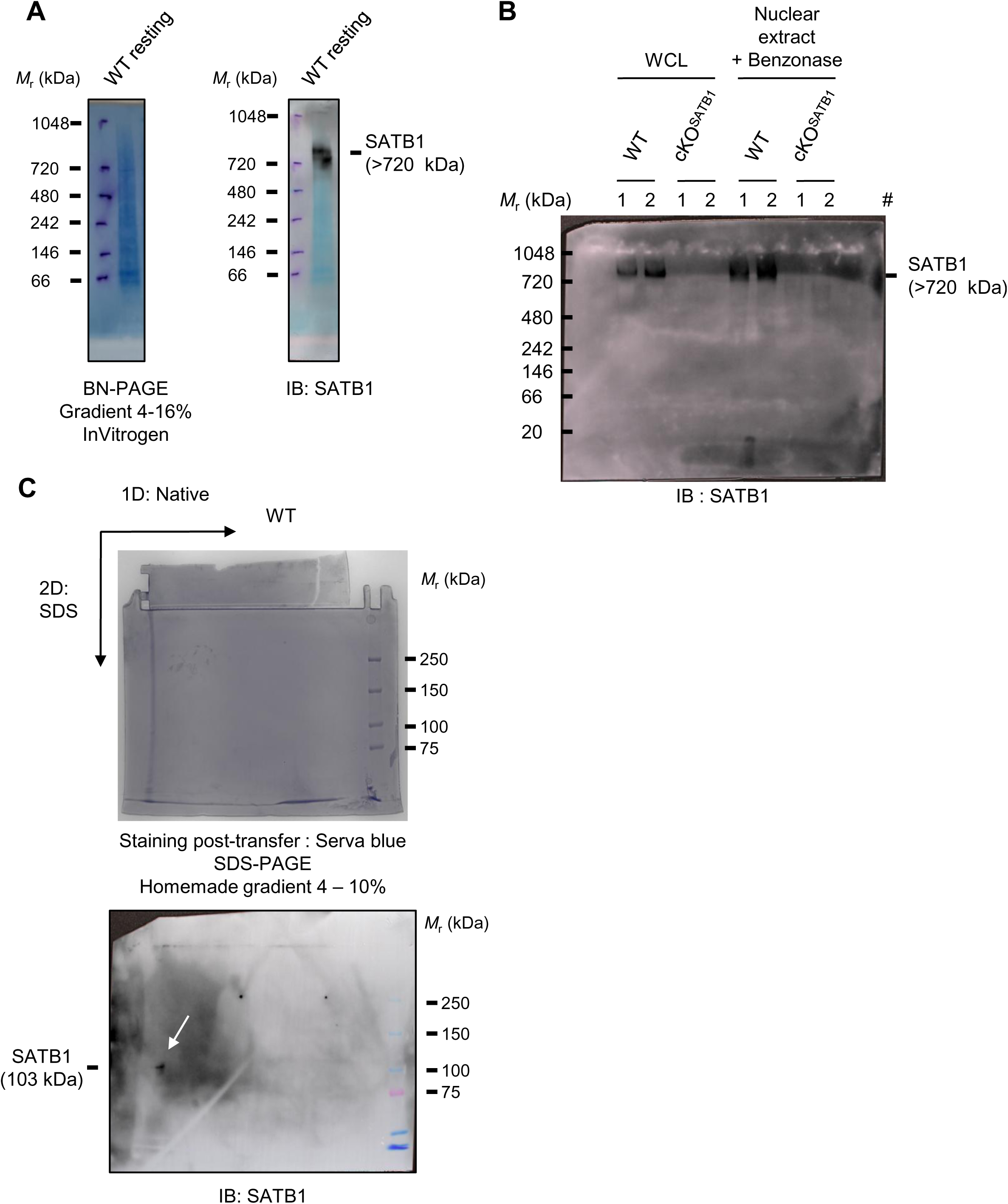
SATB1 assembles into a native >720 kDa complex in mature B-cells. **(A)** Whole-cell lysates from WT resting B-cells was resolved by blue native PAGE (BN-PAGE, 4–16% gradient, Invitrogen, BN2111BX10). The gel was stained with Coomassie G-250 to visualize total protein (left panel) and subsequently immunoblotted with anti-SATB1 antibody (1:5,000, ab109122, Abcam) (right panel). **(B)** Representative immunoblot of two independent (#) WT and cKOSATB1 whole-cell lysates (untreated) or nuclear extracts (benzonase-treated) resolved on a homemade 4–10% native gradient gel with SATB1 antibody. **(C)** Benzonase-treated nuclear extracts (10 μg) from WT resting B cells were first resolved by native PAGE (4– 16% gradient; first dimension). The entire native lane was excised and subjected to denaturing SDS-PAGE (12%; second dimension). Total protein was visualized using stain-free technology (Bio-Rad; top panel, followed by immunoblotting with anti-SATB1 antibody (bottom panel), revealing a single SATB1 spot (white arrow).

**Figure S6.**
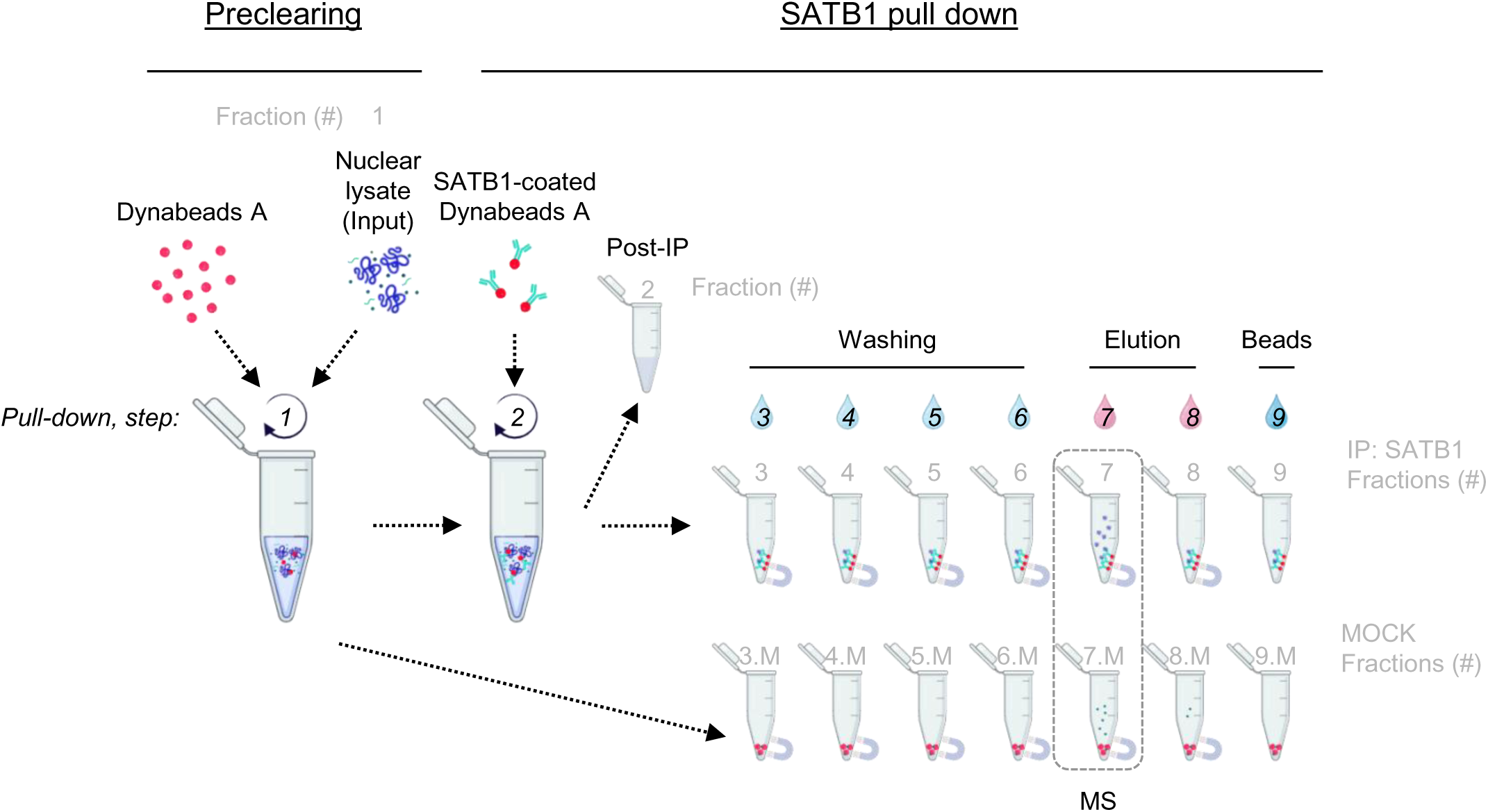
SATB1 immunoprecipitation (IP) and immunoblotting (IB) workflow. Experimental workflow of the co-immunoprecipitation mass spectrometry (co-IP/MS) approach used to detect putative SATB1 partners. Nuclear extracts were isolated from resting or LPS-activated splenic B-cells of WT or cKOSATB1 mice (aged 2–4 months) and pooled. First, 150 μg of nuclear material was incubated with Dynabeads® protein A only (MOCK). Subsequently, the pre-cleared material was incubated with beads coated with mouse anti-SATB1 antibody (sc-376096, Santa Cruz) for immunoprecipitation (IP). The IP procedure was as follows: **Step 1:** Nuclear extract (120 μl) was incubated with MOCK beads (20 μl) overnight at 4 °C. **Step 2:** The pre-cleared nuclear extract was transferred to anti-SATB1 antibody-coated Dynabeads protein A (10 μl; 0.05 μg/μl) and incubated overnight at 4 °C. **Steps 3–4:** Post-IP nuclear lysate was collected (fraction 2), and the beads were washed twice with buffer A (30 μl each; fractions 3 and 4). **Steps 5–6:** Beads were washed twice with buffer B (30 μl each; fractions 5 and 6). **Steps 7–8:** Bound proteins were eluted twice with 8M urea (20 μl then 15 μl) for 10min on ice (fractions 7 and 8). **Step 9:** Beads were resuspended in 20 μl of 1× Laemmli buffer (fraction 9). All fractions were collected. From the IP eluate (fraction 7, and fraction 7.M), 5 μl was collected for SDS-PAGE (12%) and immunoblotting, while the remainder (15 μl) was subjected to MS analysis. MOCK beads (step 1) were processed identically as SATB1-coated beads starting from step 3, and the eluate from MOCK beads is referred to fraction 7.M. MOCK beads were further resuspended in 20 μl of 1× Laemmli buffer (fraction 9.M).

**Figure S7.**
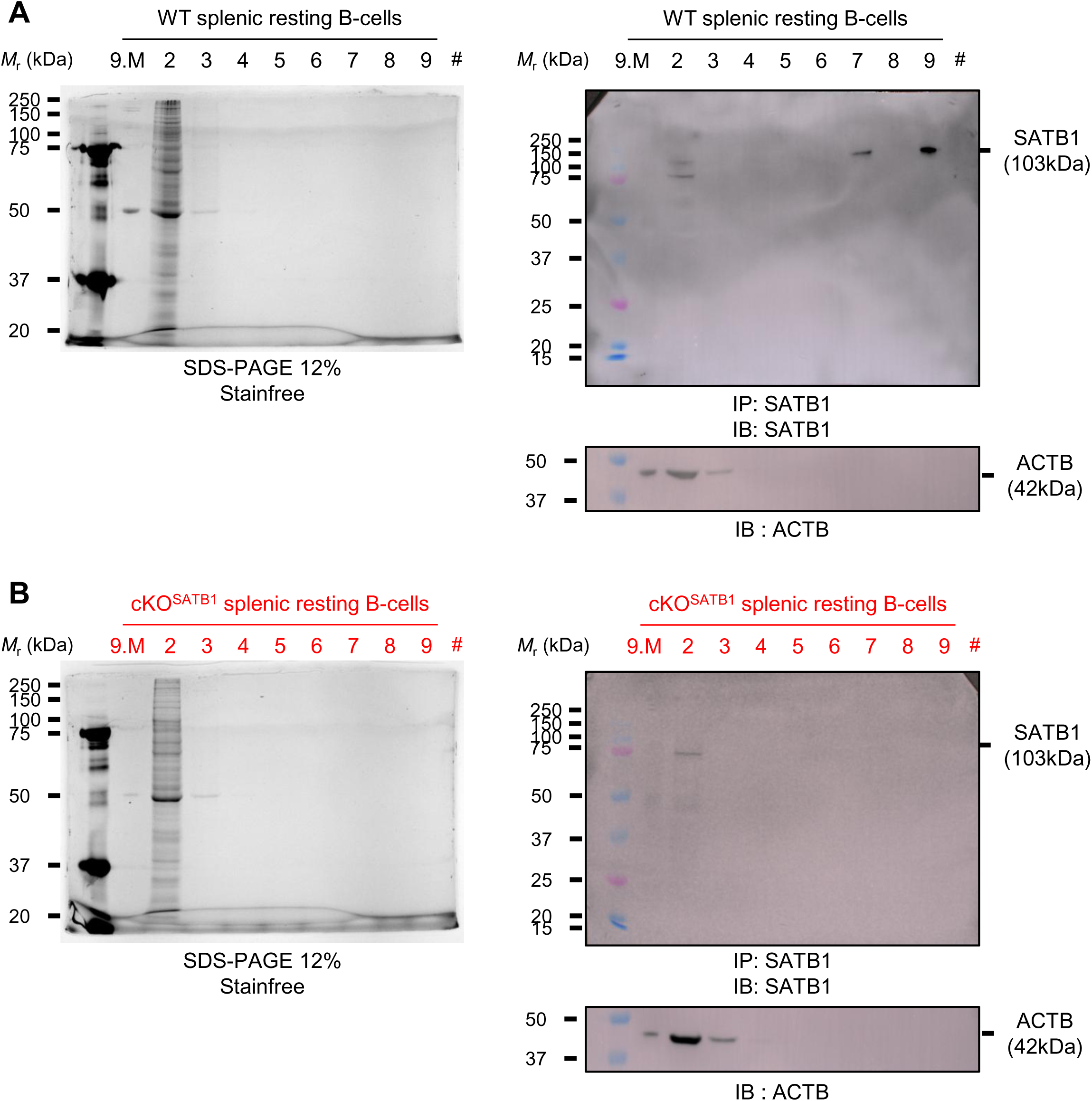
SATB1 immunoprecipitation. **(A)** Representative stain free of total protein levels and western blots with anti-SATB1 antibody (1:5,000, ab109122, Abcam) images after Immunoprecipitation (IP) performed from nuclear extracts of WT splenic resting B-cells. Fractions were collected and resolved by SDS-PAGE (12%). Fraction numbers (#) correspond to those described in Material and Methods and Figure S6. **(B)** As in *(A)* from nuclear extracts of cKOSATB1 splenic resting B-cells.

**Figure S8.**
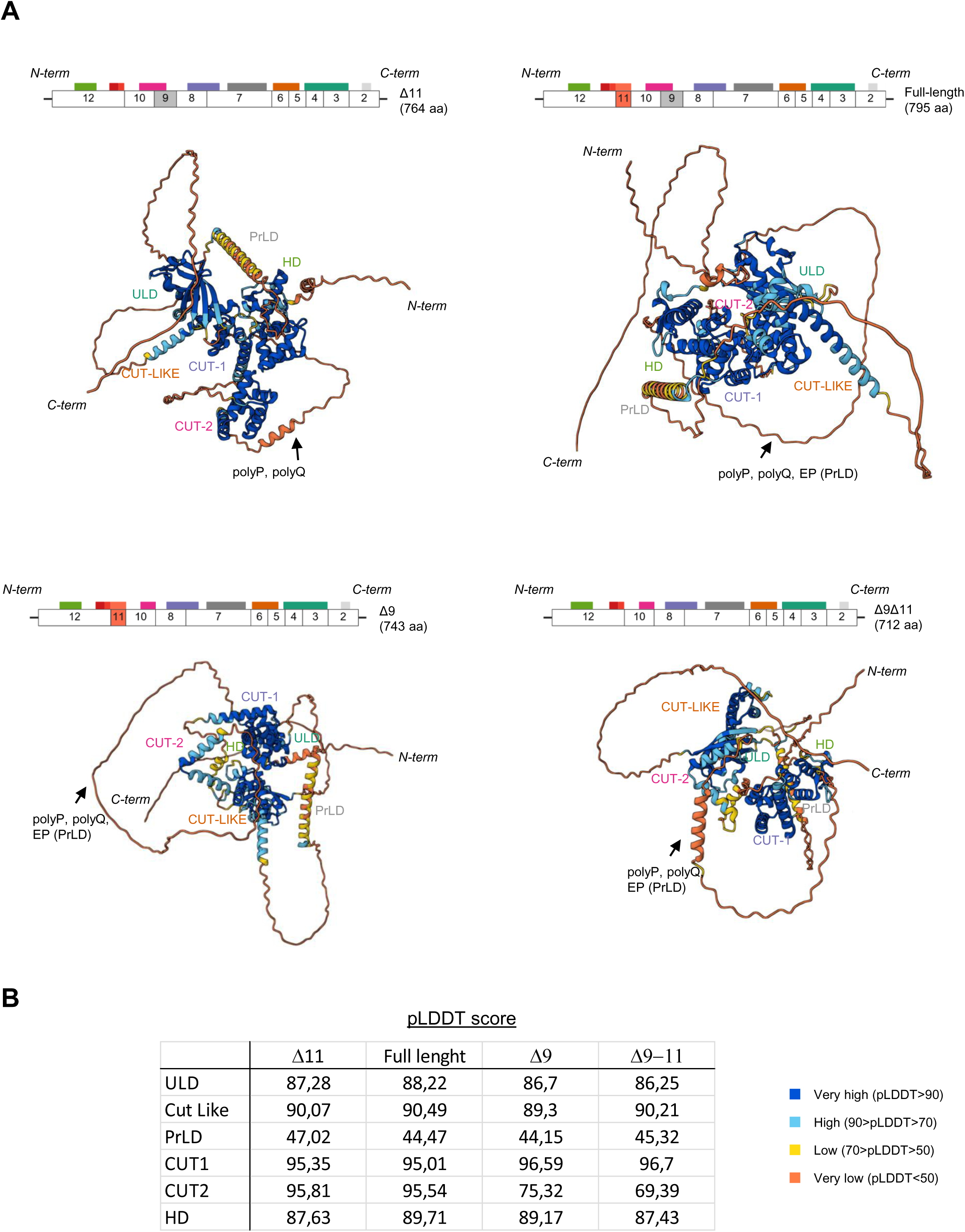
Structural prediction of murine SATB1 protein domains. **(A)** Conformation prediction of the four SATB1 isoforms using AlphaFold. **(B)** Predicted local distance difference test (pLDDT) scores for ULD, Cut Like, PrLD, CUT1, CUT2 and HD domains. The polyproline (polyP), polyglutamine (polyQ), and extra peptide (EP) domain are located by arrow

